# Network-scale disruption of RNA-protein interactions by a bioaccumulating small-molecule drug

**DOI:** 10.64898/2026.07.23.740326

**Authors:** Xiaoteng Jiang, Simone Mozzachiodi, Indra Roux, Mehul Makwana, Weiwei Zhou, Eneko Villanueva, Stephan Kamrad, Anna E. Lindell, Nonantzin Beristain-Covarrubias, Kishor D. Ingole, Nianshu Zhang, Sandip Kumar Patel, Sonja Blasche, Kathryn S. Lilley, Kiran Raosaheb Patil

## Abstract

Many small-molecule drugs accumulate in both host and gut microbial cells. Yet, the molecular consequences of intracellular drug accumulation are largely unknown. Here we show that the antidepressant duloxetine broadly disrupts RNA-protein interactions revealing a previously unrecognized mode of drug effect. Across phylogenetically diverse bacteria and human intestinal cells, duloxetine consistently reduced RNA-binding capacity, with approximately 80% of RNA-binding proteins responding to duloxetine in bacteria (*E. coli* IAI1) and human (Caco-2) cells. As a mechanistic example, we show how duloxetine disrupts interactions between the pyrimidine biosynthesis enzyme PyrB and transcripts encoding metabolically linked functions. Biochemical and structural analyses show that duloxetine competes with aspartate, the natural substrate of PyrB, weakening the enzyme’s binding with 3′ UTR-localized stem-loop structures in its RNA partners. Overall, our results uncover the off-target disruption of RNA-protein interactions due to drug accumulation and have implications for understanding molecular basis of variation in drug efficacy and toxicity.

## Introduction

Pharmaceuticals, including both antibiotics and human-targeted drugs, are a major factor underlying the dynamics and inter-individual variation in the human gut microbiota ^1–4^. These small-molecule compounds can impact the gut microbiota directly through growth inhibition ^5,6^, as well as indirectly by disrupting interactions among microbiome members and altering the host-microbiome crosstalk ^3,7–9^. These drugs can in turn be affected by the bacteria through chemical modifications (biotransformation) ^10,11^ or intra-cellular sequestration (bioaccumulation) ^5,12^, with implications for both the host and the microbiota ^13–15^. While drug biotransformation can often be attributed to specific enzymatic activities ^11,16,17^, mechanistic basis and consequences of bioaccumulation remain elusive.

Bioaccumulation can concentrate drugs hundreds of times above their extracellular levels. For example, exposure to micromolar levels of duloxetine, a widely used antidepressant, can lead to intracellular accumulation to millimolar levels, comparable to that of abundant native metabolites like amino acids ^12^. Similarly, human cells and tissues are known to markedly accumulate diverse drugs ^18–20^. Our previous work has shown that bioaccumulation impacts secretion of metabolites, affecting the growth of gut community members ^5,12^. Yet, how vital molecular processes are affected by the drug remains largely unclear, particularly since bioaccumulating bacteria often lack the canonical targets (e.g., serotonin transporters for duloxetine) and do not metabolize the compound, leaving a large gap in our understanding of drug mode of action and off-target effects. This knowledge gap also extends to human cells wherein duloxetine exhibits significant pharmacological promiscuity, binding to unintended off-target proteins and metabolic enzymes that complicate its systemic interaction networks and side-effect profile ^21,22^.

We previously noted that bioaccumulated duloxetine preferentially alters the thermal stability of bacterial metabolic enzymes, which are enriched in nucleotide cofactor-binding domains, particularly the widely conserved Rossmann fold ^12^. In parallel, enzymes harboring Rossmann fold have been identified as RNA-binding proteins (RBPs) across diverse species, a dual-function phenomenon termed “moonlighting” ^23–25^. In these moonlighting enzymes, the RNA-binding interfaces are frequently coupled to the catalytic pockets either through direct spatial overlap or dynamic allosteric regulation, creating vital riboregulatory nodes at the interface of metabolism and post-transcriptional control ^26–29^. Since RBPs function as post-transcriptional hubs coordinating rapid metabolic adaptations to environmental changes ^30–33^, the disruption of RNA-protein binding could have consequences for metabolic activity and cellular growth. We therefore hypothesize that intracellular accumulation of duloxetine could compete for these conserved structural folds in RBPs and systemically perturb riboregulation.

To test our hypothesis, we here investigate the impact of duloxetine bioaccumulation on RNA-binding proteins by employing orthogonal organic phase separation combined with data-independent acquisition (OOPS-DIA) mass spectrometry ^25,34^. We thus uncovered a network-wide disruption of RNA-binding capacity as a consequence of drug accumulation, impacting, among others, metabolic enzymes conserved across kingdoms.

Pyrimidine biosynthesis enzyme PyrB emerged as a novel and highly conserved RBP. Multi-omics and structural analyses reveal that duloxetine does not merely act as a classical competitive inhibitor at the catalytic pocket; rather, its binding triggers a conformational disruption of the enzyme’s distinct RNA-binding interface forming a hitherto unknown riboregulatory circuit in nucleotide biosynthesis. Our data thus unveils a novel, cross-kingdom mechanism: disruption of riboregulatory networks by a small molecule drug.

## Results

### Duloxetine bioaccumulation remodels the bacterial RNA-binding proteome and targets essential metabolic enzymes

To understand the mechanistic link between the antidepressant duloxetine bioaccumulation and microbiome physiology, we started by evaluating the impact in growth of four phylogenetically distinct bioaccumulating gut bacteria (*Escherichia coli* IAI1, *Bacteroides uniformis*, *Lacticaseibacillus paracasei*, and *Streptococcus salivarius*) and a non-bioaccumulating control strain, *E. coli* ED1a. We used a physiologically relevant concentration of duloxetine (50 μM) and modified Gifu Anaerobic Medium (mGAM), conditions consistent with previously established drug accumulation phenotypes, enabling direct comparison ^6,12^. Compared to growth with the vehicle control (0.1% DMSO), duloxetine induced a decline in the optical density of *E. coli* IAI1 during the stationary phase and caused log-phase changes in *B. uniformis* and *L. paracasei*, while it had no effect on the non-bioaccumulating *E. coli* ED1a (Figure S1A).

Since duloxetine-interacting proteins are enriched in conserved Rossmann-fold metabolic enzymes with moonlighting RNA-binding potential ^12,24,25^, we reasoned that duloxetine-induced growth inhibition may involve disruption of RNA-protein regulatory networks. We therefore applied orthogonal organic phase separation (OOPS) method for simultaneous enrichment and quantification of RNA-bound proteins and their associated RNAs from crosslinked RNA-protein complexes under duloxetine exposure (Figure 1A).

**Figure 1.**
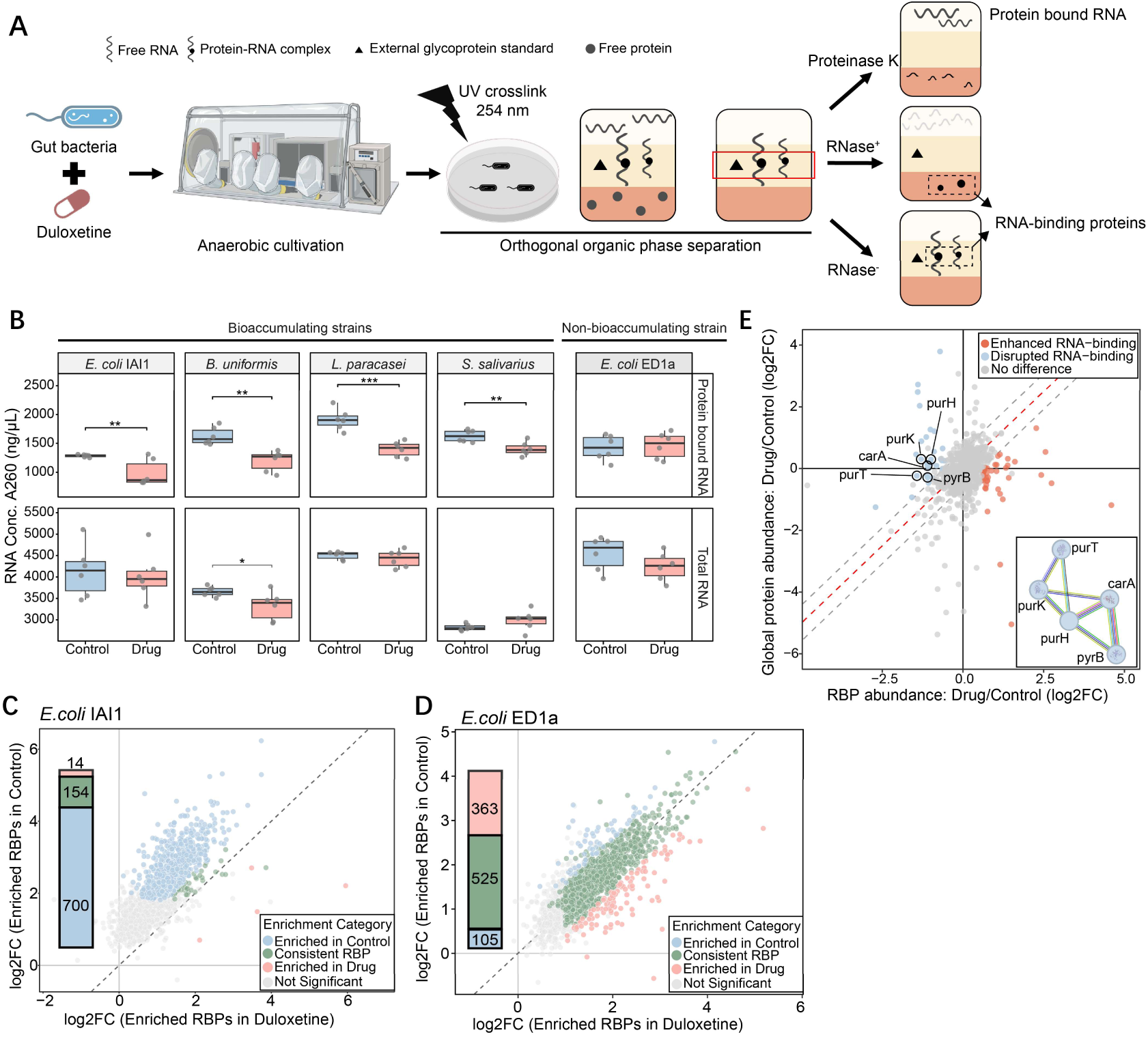
Duloxetine bioaccumulation disrupts RNA-protein interactions of *E. coli* IAI1 with *de novo* UMP/IMP biosynthesis process emerging as a key network. **A.** Duloxetine treatment of gut bacteria and RNA-protein complexes characterization workflow. Bacteria treated with duloxetine or DMSO control were cultivated anaerobically, with growth monitored hourly until reaching the stationary phase. Following UV crosslinking at 254 nm, bacteria were lysated and underwent an initial orthogonal organic phase separation (OOPS) to isolate crosslinked RNA-protein complexes at the interphase. For RNA testing, the interphase was treated with Proteinase K. To identify RNA-binding proteins, the interphase fraction was divided and treated with or without RNase (RNase^+^/RNase^-^), followed by a second phase separation and analyzed via LC– MS/MS in data-independent acquisition (DIA) mode. Proteins released from the interphase exclusively upon RNase treatment (RNase-sensitive proteins) were RNA-binding proteins, defined by their significant depletion in the RNase⁺ fraction relative to the RNase⁻ control. **B.** RNA recovered from RNA-bound proteins (top) and total RNA (bottom) among bioaccumulating and non-bioaccumulating bacteria, *P < 0.05, **P < 0.01 (two-sided Student’s t-test). **C.** Comparison of significantly enriched RNA-binding proteins (RBPs) with and without duloxetine treatment in *E. coli* IAI1. The scatter plot displays the RNA-dependent enrichment (log₂ fold change of RNase-untreated vs. RNase-treated samples) of each quantified protein under duloxetine treatment (X-axis) versus control conditions (Y-axis). The dashed diagonal line represents the line of y = x, indicating unchanged RNA-binding capacity under the two conditions. Each dot represents a quantified protein detected in both conditions. Significantly enriched RBPs were defined by FDR-adjusted P < 0.01 and log₂ fold change > 1. The inset bar plot summarizes the total number of enriched RBPs in each condition, including proteins that were completely depleted upon RNase treatment across all replicates. These proteins are considered high-confidence RBPs due to their extreme RNase sensitivity but are not represented in the scatter plot because of missing values in quantitative measurements. **D.** Comparison of significantly enriched RBPs with and without duloxetine treatment in the non-bioaccumulating strain *E. coli* ED1a. Plot layouts, axis definitions, and significance thresholds are identical to those described in (**C**). **E.** Four-quarter plot distinguishing RNA-binding alterations from global protein expression changes. The scatter plot shows the Log_2_ fold change of RBP abundance (x-axis) versus global protein abundance (y-axis) under duloxetine treatment compared to the control. The red dash line depicts the 1:1 trend (y = x) of the whole proteome and RBP changes. Proteins within the gray dash line indicate the abundant change as an RBP corresponds to its protein expression level (a difference boundary within ± 0.58 log_2_ fold change). Proteins represented by coloured dots are those whose changes in RBP abundance do not correspond to changes in their overall expression, where red corresponds to increased RNA-binding and blue to decreased RNA-binding (defined by FDR-adjusted P < 0.05, absolute log2 fold change in RBP abundance > 0.58). Inset, the duloxetine-disrupted RBPs corresponding to the *de novo* UMP/IMP biosynthetic process analyzed by STRING.

In all four bioaccumulating strains, duloxetine treatment significantly reduced the yield of protein-bound RNA, despite total RNA levels remaining largely unchanged (Figure 1B). This indicates a widespread disruption of the RNA-protein interactome by duloxetine. In contrast, the non-bioaccumulating strain *E. coli* ED1a showed no such reduction, suggesting that the effect in the other strains is due to bioaccumulation (Figure 1B).

To characterize the affected RNA-binding proteins, we focused our deep proteomic profiling on the two gut *E. coli* strains (IAI1 and ED1a). This comparison controls for general drug exposure to pinpoint bioaccumulation-specific effects, and benefits from the most comprehensively annotated bacterial proteome. Given that the catalog of gut bacterial RNA-binding proteins (RBPs) is expanding as a new frontier in microbiology with many RBPs remaining incompletely annotated ^32,35–38^, we first validated that our OOPS-DIA method robustly captures known RBPs as well as previously unannotated RBP candidate. By quantifying changes in protein abundance upon RNase treatment and applying stringent thresholds (log2FC > 1, adj. P < 0.01), we identified 993 and 868 high-confidence RBPs in *E. coli* ED1a and IAI1, respectively (Figure S1B and S1C, Supplementary TableS1). The specificity of our workflow was validated via comprehensive functional enrichment: Gene Set Enrichment Analysis (GSEA) highlighted a strong enrichment signature for the RNA binding GO term (GO:0003723) (Normalized Enrichment Score > 2.7) (Figure S1D and S1E), while KEGG and InterPro analyses enriched for canonical RNA-related functions and domains such as RNA metabolism and modification (Figure S1F-S1I and Figure S2A-S2B).

At the proteome level, consistent with the broad reduction of protein-bound RNA described above, we found a drastic remodeling of RNA-binding proteins in the bioaccumulating strain IAI1. Duloxetine treatment reduced the number of significantly enriched RBPs from 854 to 168, representing an 80% decrease (Figure 1C). This extensive loss is captured by the RNA-binding proteome shifting strongly toward the control Y-axis, away from the y = x diagonal, suggesting that duloxetine broadly abolished their RNA-dependent enrichment. Such remodeling was absent in the non-bioaccumulating ED1a strain (Figure 1D), highlighting that drug bioaccumulation is essential for this RNA-protein disruption.

To decouple binding disruption from general reduction in protein abundance, we cross-referenced the RNA-binding proteome dynamics with global proteome changes in IAI1. We identified 36 RBPs whose RNA association was significantly reduced without corresponding changes in their abundance (Figure 1E). This indicates that duloxetine disrupts the RNA-binding activity of these proteins at the post-translational level, independent of their cellular stability. Mapping the physical and functional interaction networks of these 36 RBPs by STRING ^39^ revealed three functionally interacting clusters: (i) *de novo* UMP/IMP biosynthetic process, (ii) glyoxylate and dicarboxylate metabolism, and (iii) phosphatidylglycerol metabolic process, which is linked to signaling and cell division (Figure S2C). The RBPs disrupted by duloxetine are positioned at the critical nodes of *de novo* nucleotide biosynthesis spanning both the pyrimidine (CarA, PyrB) and purine (PurT, PurK, PurH) branches (Figure S2D), which suggests the drug may perturb nucleotide biosynthesis. Notably, the convergence of our OOPS-DIA data with a previous thermal-proteome profiling dataset ^12^ identifying drug-binding targets (Figure S2C), strongly suggests a direct interaction mechanism: duloxetine binds to these novel moonlighting RBPs and physically disrupts their ability to interact with RNA.

### The novel moonlighting RBP PyrB drives a riboregulatory feedback loop vulnerable to duloxetine

To understand how duloxetine interacts with the RBPs, we conducted molecular docking simulations using duloxetine as the ligand against their AlphaFold-predicted structures. All candidates from the identified *de novo* nucleotide biosynthesis pathway exhibited strong binding affinities (scores < −6 kcal/mol), while five non-target proteins randomly drawn from our unresponsive OOPS-DIA background showed weaker binding (average scores > −5.4 kcal/mol) (Figure S2E). Among them, PyrB, the conserved catalytic subunit of aspartate transcarbamoylase that governs the rate-limiting step of pyrimidine biosynthesis ^40^, showed the highest predicted affinity. Notably, the CATH (Class, Architecture, Topology, Homology) database ^41^ verifies that PyrB contains a canonical Rossmann fold, providing a structural basis for its drug susceptibility. Indeed, our docking models predict that duloxetine forms hydrogen bonds, Pi-cation, and Pi-Alkyl interactions with PyrB (Figure S2F). Given its essential rate-limiting metabolic function and perfect structural alignment with our proteome-wide binding patterns, PyrB emerged as the primary mechanistic target driving the observed growth inhibition

To characterize the foundational RNA-binding landscape of the novel RBP – PyrB, we first validated its interacting with RNA *in vitro* using electromobility shift assays (EMSA) with total RNA from *E. coli* IAI1. Our results suggest that PyrB forms RNA-protein complexes, whereas the addition of duloxetine effectively hinders complex formation, accompanied by drug-induced protein precipitation at high concentrations (Figure 2A, S3A, and S3B). We then performed individual-nucleotide resolution UV crosslinking and immunoprecipitation (iCLIP) to capture the exact RNA interactome *in vivo*. The results confirmed PyrB as a *bona fide* RBP, supported by the RNase sensitivity of the immunoprecipitated RNA-PyrB complexes (Figure 2B and S3C). Consistent with our OOPS-DIA data, duloxetine treatment reduces the overall RNA-PyrB complex signal intensity (Figure 2B), confirming the drug’s capacity to disrupt the RNA-PyrB interactions *in vivo*. Analysis of these specific targets showed that PyrB preferentially interacts with intergenic and protein-coding regions via distinct motifs (Figure S3D-S3F, S3L). Beyond linear sequence recognition, Positionally enriched k-mer analysis (PEKA) showed downstream GC-rich enrichments (Figure S3G) alongside characteristic positional patterns, notably a dual-peak GUUCUC cluster (Figure S3H), suggesting local RNA secondary structure features may contribute to PyrB recognition.

**Figure 2.**
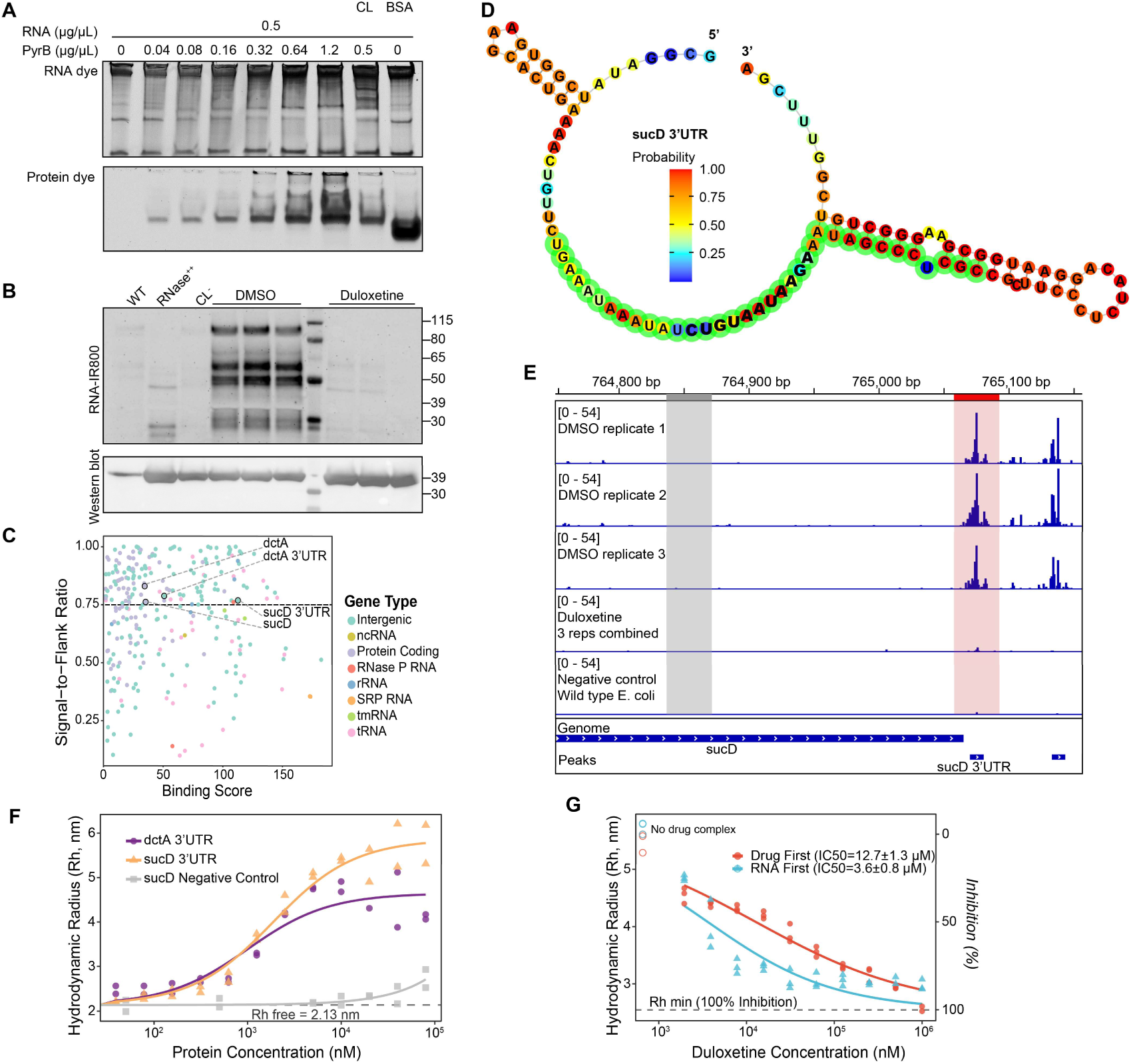
PyrB is a novel RBP and duloxetine disrupts its RNA interactions. **A.** Electromobility shift assay (EMSA) of PyrB and the whole RNA extracted from the IAI1 strain. The gel was stained with SYBR Green first to show the RNA pattern and later with SYPRO Ruby on the same gel to show the protein position. The protein smear suggests the PyrB-RNA complex. CL, UV cross-linked PyrB-RNA complex. BSA, Bovine Serum Albumin, is used as a negative control in this experiment (Protein concentration: 1.6 µg/µL). **B.** Visualization of biotin-labeled RNA (upper) and PyrB (bottom) on the same nitrocellulose membrane. WT, wild type IAI1. RNase^++^, PyrB overexpression sample treated with excessive amount of RNase. CL^-^, PyrB overexpression sample without UV cross-link. DMSO, PyrB overexpression samples with 0.1% DMSO. Duloxetine, PyrB overexpression samples treated with 50 μM duloxetine. **C.** 2D score plot of the iCLIP peaks. The y-axis shows signal to flank ratio. High score suggests high specificity of RNA-protein binding. X-axis shows binding scores as calculated by PureCLIP, indicating statistical confidence. **D.** RNA structure of the PyrB target *sucD* RNA 3’UTR predicted by ViennaRNA. The green highlighted regions (35 nt) were expanded from the 11 nt PyrB-interacting RNA regions. Enlarged nucleotide letters represent the 11 nt binding sites. **E.** iCLIP signal tracks of the DMSO control (3 replicates), the merged signal of duloxetine treatment (from 3 replicates), and the wild type *E. coli* negative control of the *sucD* RNA. The highlighted red/grey regions (35 nt) represent the PyrB interacting area and the non-interacting area, respectively. These regions were selected for RNA synthesis and validation. The bottom Genome/Peaks track shows the coding region of *sucD* and the PureCLIP identified 11nt binding regions of PyrB. **F.** Microfluidic diffusional sizing (MDS) binding profiles of PyrB with target RNAs. Hydrodynamic radius measurements were performed using 20 nM of 5’ Cy5-labeled RNA probes (excitation at 647 nm) corresponding to the *dctA* 3’ UTR, *sucD* 3’ UTR, and a structurally dissimilar *sucD* negative control. Data were fitted using a non-linear least squares (nlsLM) regression model (two technical replicates). **G.** Competition binding assays measured by MDS reveal the displacement of 25 nM *sucD* 3’ UTR RNA by increasing concentrations of duloxetine. The left Y-axis represents the absolute hydrodynamic radius, while the right Y-axis indicates the normalized inhibition percentage. Drug first: Pre-incubation strategy. PyrB was first incubated with the duloxetine gradient prior to RNA addition. RNA first: Displacement strategy. PyrB was pre-assembled with RNA to form the complex before duloxetine titration. Solid lines represent a non-linear logistic regression fit (2 parameters) with locked R_h_complex_ (0 µM duloxetine, no drug complex) and R_h_min_ (BSA + RNA control) baselines (three technical replicates).

Indeed, among the highly specific iCLIP targets (signal-to-flank ratio > 0.75, binding score > 30), PyrB crosslink sites were consistently located within regions connecting two predicted stem-loop structures in both the protein-coding sequences and the 3’ UTRs of the *dctA* and *sucD* mRNAs (Figure 2C, 2D, and S3I-S3K). Strikingly, both genes are linked to PyrB’s core metabolic function. DctA imports L-aspartate, the direct substrate for PyrB ^42^. Meanwhile, SucD generates ATP and succinate; ATP acts as an allosteric activator of this pyrimidine pathway, while succinate accumulation directly occupies PyrB’s catalytic pocket to inhibit its activity ^43,44^. This profound metabolic convergence—where an enzyme binds to the mRNAs encoding its own substrate-provider and allosteric-regulator—strongly suggests a sophisticated and coordinated riboregulatory feedback loop. Crucially, the iCLIP signals on the 3’ UTRs of both *dctA* and *sucD* were diminished by duloxetine (Figure 2E, S4A, and S4B), proving that the drug mechanically disrupts this regulatory network.

To quantitatively validate these structure-based RNA-PyrB interactions, we performed microfluidic diffusional sizing measurements using synthesized 35-nucleotide RNA fragments identical to the 3’ UTR binding regions of *dctA* and *sucD* transcripts (Figure 2D and S3J,). PyrB exhibited high-affinity binding to these structured targets, yielding tight dissociation constants (K_D_) of 0.99 ± 0.28 µM and 1.87 ± 0.28 µM, respectively (Figure 2F). In contrast, a structurally dissimilar sequence derived from an upstream region of *sucD* 3’ UTR showed no measurable interaction with PyrB, further confirming the structural specificity of PyrB-RNA recognition (Figure 2E, 2F and S4C).

Finally, we investigated the thermodynamic interplay between RNA binding and drug inhibition using microfluidic diffusional sizing. While duloxetine effectively disrupted the PyrB-RNA complex regardless of the order of RNA/drug addition, the system exhibited remarkable allosteric behavior: the protein’s sensitivity to the drug varied depending on its initial RNA-bound state (Figure 2G). Displacing RNA from a pre-formed PyrB-RNA complex yielded a highly potent IC50 of 3.60 ± 0.83 µM (Hill coefficient 0.60). In contrast, preventing RNA from binding to a pre-incubated PyrB-drug complex resulted in a reduced inhibitory potency, shifting the IC50 to 12.67 ± 1.32 µM (Hill coefficient 0.47). This varied sensitivity indicates that RNA binding shifts PyrB into a conformation that is more susceptible to duloxetine’s disruptive insertion, providing initial yet compelling evidence of allosteric modulation.

### Duloxetine occupies the enzyme pocket of PyrB, causing global structural changes

To understand the structural basis of duloxetine-induced RNA disassociation, we modeled its interaction with PyrB. PyrB has two major conformations: the active state (PDB 4kgv) and the inactive state (PDB 1r0c) ^45,46^. Molecular docking simulations showed that in the active state, duloxetine competitively occupies the catalytic pocket near its Rossmann-fold structure (molecular docking score, MDS = −7.38 kcal/mol), displacing its natural substrate, aspartate (MDS = −5.7 kcal/mol) (Figure 3A). Detailed interaction analysis identified three key hydrogen bonds: the amine hydrogen (H7) of duloxetine interacts with the carbonyl oxygen of Ala52 (2.0 Å), while its thiophene sulfur (S1) and ether oxygen (O1) form stable hydrogen bonds with the guanidinium groups of Arg106 (2.79 Å) and Arg230 (2.65 Å), respectively (Figure 3A). Notably, the sulfur-containing thiophene ring of duloxetine precisely overlaps with the O2, O4, and C2 positions of the substrate aspartate, effectively hijacking the critical anchor points (Arg106 and Arg230) required for substrate stabilization ^47^. Meanwhile, the bulky naphthalene ring remains positioned toward the outer rim of the catalytic pocket, physically blocking the entry path for aspartate.

**Figure 3.**
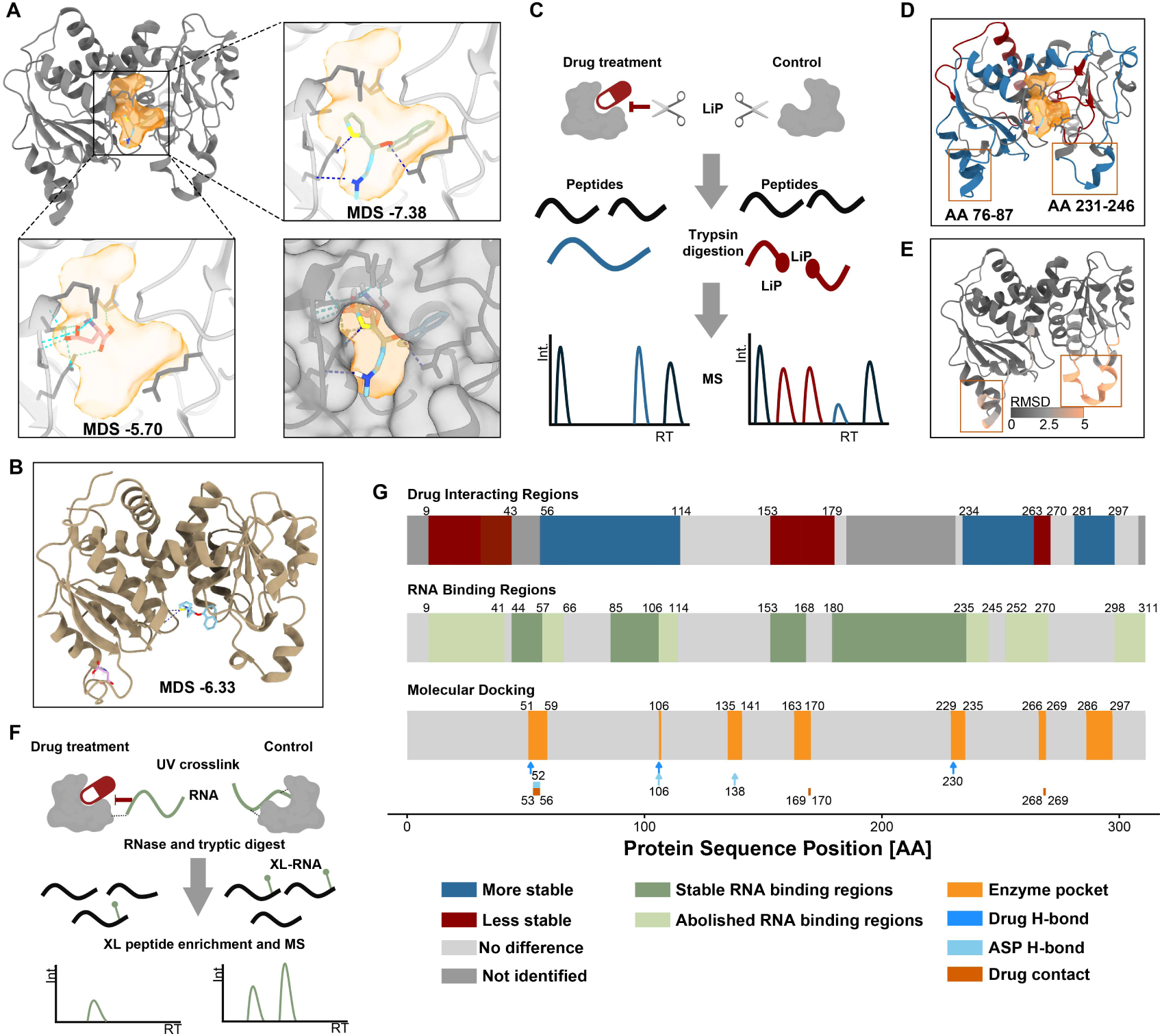
Duloxetine binds PyrB in its enzyme pocket and rearranges its conformation. **A.** Molecular docking of duloxetine (zoom-in at top right) and the natural substrate aspartate (zoom-in at bottom left) within the active state of PyrB (PDB: 4kgv). The enzyme’s catalytic pocket is highlighted with an orange surface. The bottom-right panel displays the structural overlay of duloxetine and aspartate, illustrating direct steric competition for the active site. The semi-transparent grey surface representation in this panel demonstrates how the bulky moiety of duloxetine physically blocks the substrate entry regions. MDS: molecular docking score. **B.** Molecular docking of the inactive state PyrB (PDB 1r0c) with duloxetine (light blue) and aspartate (pink). **C.** Schematic workflow of LiP-MS. Duloxetine-binding changes the accessibility of LiP digestion, resulting in different peptide patterns. The blue peptide suggests structurally more stable region (less accessible to LiP). The red peptides suggest structurally less stable regions (more accessible to LiP). **D.** Visualization of the structurally changed peptides mapped against the structure of PyrB. Peptides generated from the more stable structure after binding with duloxetine are colored in blue, suggesting more proteolytic resistant. Peptides generated from the less stable structure after binding with duloxetine are colored in red, suggesting less proteolytic resistant. Peptides in the squares suggest the regions with most distinct structures between the active and inactive state PyrB. The enzyme active site pocket is highlighted in orange. **E.** The structural similarity of the active and inactive state PyrB (representing model: active state of PyrB, PDB 4kgv). Color range from grey to pink denotes the Root Mean Square Deviation (RMSD) score. High score indicates a high degree of dissimilarity. The two orange squares indicate the most distinct regions between the two states of PyrB. **F.** Schematic workflow of the RNA-XL-MS. Samples treated with duloxetine and DMSO control were UV crosslinked, RNase and trypsin digested, RNA-XL-peptides enriched, and mass spectrometry analysis. **G.** Multi-track sequence projection of PyrB. The top track displays duloxetine-induced structural alterations identified by LiP-MS (blue: increased stability after drug binding; red: decreased stability; grey: no significant difference). The middle track illustrates the RNA interactome mapped by RNA-XL-MS, distinguishing between stable (dark green) and abolished (light green) RNA-binding regions following drug treatment. The bottom track annotates the predicted enzyme pocket (orange) and specific atomic interactions, including essential hydrogen bonds (blue arrows for duloxetine; light blue for aspartate). Drug contact (dark orange) is defined as residues within a <4.0 Å distance threshold from duloxetine using ChimeraX. Numbers indicate amino acid sequence positions.

In the inactive state, the catalytic pocket adopts a low-affinity conformation that fails to stably bind aspartate. However, our docking simulation reveals that this open conformation can accommodate the bulkier duloxetine molecule (MDS = −6.33 kcal/mol, maintaining 2 hydrogen bonds) (Figure 3B). By occupying this space, we hypothesize that duloxetine induces conformational changes in PyrB, locking the enzyme and preventing catalytic activation.

To capture the structural changes induced by duloxetine, we applied limited proteolysis-mass spectrometry (LiP-MS) ^48^ to locate the duloxetine binding sites of PyrB. Based on the different resistance to proteolytic activity upon duloxetine treatment, as indicated by the distribution and the intensity of fully tryptic peptides (more proteolytic resistant, more stable structure), and semi-tryptic peptides (less proteolytic resistant, less stable structure), we located the peptides from regions undergoing structural change upon duloxetine binding (Figure 3C, 3D and S4D). Peptides corresponding to the predicted duloxetine binding site and the substrate entry showed significant protection from proteolytic activity upon duloxetine treatment. Strikingly, key loops that move together from the inactive state to the active state (AA76-87, 231-246) were stabilized after duloxetine binding (Figure 3D, 3E and S4E). Since they are the substrate entry regions of PyrB ^49^, our result indicates that this stabilization locks PyrB in an active-like conformation, and thus prevents its substrate aspartate from entering the catalytic pocket.

Further, to determine how this catalytic pocket occupation impacts the RNA interactome, we applied the UV cross-linking mass-spectrometry method (RNA-XL-MS) ^50^ on duloxetine-treated samples and controls (Figure 3F). This method allowed us to enrich the RNA crosslinked peptides and identify the RNA-peptide interactions at amino acid resolution by directly measuring the mass shifts corresponding to specific nucleotide adducts (e.g., residual mono- or di-nucleotides) crosslinked to peptides. Our results revealed a significant drug-induced remodeling of the RNA-binding interface. In the DMSO control, we identified 20 RNA-crosslinked peptides, 17 of which overlap with regions identified by LiP-MS as conformationally sensitive. Following duloxetine treatment, the number of detectable crosslinked peptides was reduced to 10 (Figure 3G and S4G).

By projecting these RNA-XL-MS results alongside our LiP-MS and docking data onto a multi-track sequence map, we observed a high spatial overlap between drug binding, structural remodeling, and RNA dissociation (Figure 3G). We found that this loss of RNA contact was not random but reflected the drug-induced structural disruptions. For instance, the abolishment of RNA binding at the N-terminus (AA9-41) directly aligned with its structural destabilization (AA9-43). Many were overlapped or closed to the drug-stabilized enzyme pocket (AA57-66, 106-114, 235-245, 252-270, 298-311). Notably, one displaced RNA-binding peptide (AA235-245) overlaps directly with the key allosteric loop (AA231-246) structurally differentiating the active/inactive states right after the critical enzyme pocket harboring Arg230 hydrogen-bond anchor for the drug ^45^ (Figure 3D and 3G). This targeted disruption of RNA binding at the allosteric and catalytic hub suggests duloxetine disrupts the precise conformational dynamics required for RNA recognition.

### Duloxetine allosterically disrupts the RNA-binding interface

To mechanistically define the duloxetine-induced RNA displacement, we constructed a 3D model of the PyrB-RNA complex using a 100-nt sequence encompassing the 3’ UTR target in *sucD*, guided by the interaction sites identified from our RNA-XL-MS and iCLIP data.

In the absence of duloxetine, the *sucD* 3’ UTR binds to a positively charged cleft located on the posterior surface of the protein’s catalytic pocket (Figure 4A, 4B and S4F). RNA-XL-MS identified RNA-binding peptides situated at two sides of the protein, forming a clamp-like structure that anchors the 11-nt binding motif of RNA identified *in vivo* by iCLIP (Figure 4C). Upon duloxetine binding within the catalytic pocket, as our LiP-MS data suggested, PyrB is locked in an active-like, stable conformation, which draws residues 76-87 and 231-246 closer together, distorting the geometry of the distal RNA-binding cleft (Figure 4B, 4D and S4F). This conformational rearrangement is accompanied by a redistribution of electrostatic potential: negatively charged residues (e.g., Asp19 and Asp237) are introduced into the interface, while key positively charged residues (e.g., Lys8) are displaced away from the RNA-binding surface. As a result, this creates a strong electrostatic repulsion against the negatively charged RNA backbone, forcing the iCLIP-identified 11 nt binding motif to turn away from the interaction face, and leading to a loss of stable RNA engagement (Figure 4D and S4F).

**Figure 4.**
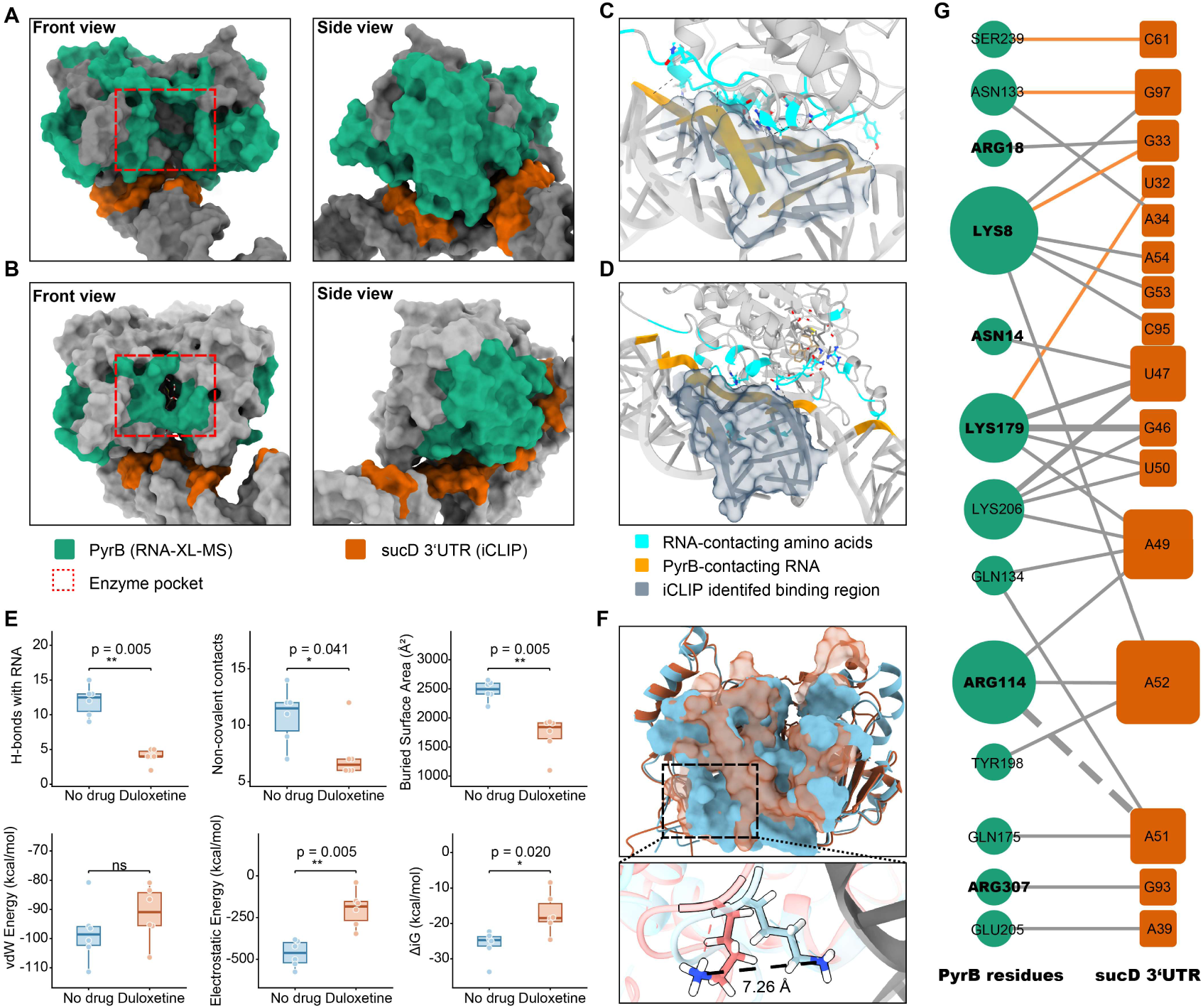
Duloxetine-binding weakens the PyrB-RNA interaction. **A, B.** Structural models of the PyrB-sucD 3’UTR complex generated by HADDOCK, in the absence (**A**) and presence (**B**) of duloxetine. Green surface indicates PyrB RNA-binding peptides identified via RNA-XL-MS. Orange surface represents RNA sequences interacting with PyrB identified by iCLIP. Left, the front view of the complex. The red dashed box highlighting the catalytic pocket (which overlaps with the natural substrate-binding site). Right panels show the side view, illustrating the protein-RNA interface. **C, D.** The detailed interaction interfaces of the PyrB-sucD 3’UTR complex. The RNA-contacting amino acids are colored in cyan (from the top 3 best-scored models). The black dashed lines are hydrogen bonds. The protein-contacting RNA is colored yellow. Duloxetine in **D** is shown in beige. The transparent dark blue area represents the 11 nt binding sites identified by iCLIP. For visual simplicity, only the best-scoring model in each state is shown. **E.** Quantitative comparison of the PyrB-sucD 3’UTR binding interface between the no duloxetine model (blue, n = 6) and duloxetine-bound model (red, n = 6) ensembles. The panels show the number of hydrogen bonds (top left), number of non-covalent contacts (top middle), buried surface area (BSA, Å²; top right), van der Waals energy (kcal/mol; bottom left), electrostatic energy (kcal/mol; bottom middle), and binding free energy (ΔiG, kcal/mol). Each box represents the median and interquartile range (IQR); whiskers extend to 1.5× IQR; individual data points correspond to single structural models. P-values were calculated by Wilcoxon ran-sum test (two-sided). *p < 0.05, **p < 0.01, ns, not significant. **F.** Buried surface area of PyrB-sucD 3’UTR interactions between the no duloxetine model (the best-scored model ingroup, blue) and the duloxetine-bound model (the best-scored model ingroup, red) shown on the aligned protein structures. Zoomed-in figure shows the special replacement of a key RNA-interacting amino acid, Lys8, away from the interacting RNA (Grey). **G.** 2D interaction network between PyrB (green circles) and the *sucD* 3’UTR (orange squares). Only hydrogen bonds predicted in at least two HADDOCK models are shown. The solid grey lines represent lost hydrogen bonds after duloxetine binding. Dashed line represents the weakened interaction after duloxetine binding (increased bond distance). The orange lines represent the newly generated hydrogen bonds after duloxetine binding. The thickness of the line represents the number of models processing hydrogen bond interactions at the current site.

To quantify the impact of the conformational rearrangement on RNA binding, we evaluated the thermodynamic binding affinities of the top models from both states. Compared to the no-drug model, the duloxetine-bound state exhibits a marked reduction in interfacial stability, with hydrogen bonds decreasing from 12.0 ± 2.2 to 4.0 ± 1.1, atomic contacts from 10.8 ± 2.5 to 7.3 ± 2.3, and buried surface area from 2474.2 ± 168.8 Å² to 1704.6 ± 322.4 Å². Electrostatic binding energy is substantially weakened (−466 ± 72 to −198 ± 99 kcal/mol), resulting in a significantly reduced binding free energy (ΔiG) (−26.0 ± 3.7 to −17.1 ± 5.2 kcal/mol) (Figure 4E). At the residue level, the RNA-binding disruption is driven by spatial displacement and side-chain reorientation of key anchoring residues. Notably, Lys8 undergoes a pronounced side-chain reorientation of 85.4°, with its terminal NZ atom shifting 7.26 Å away from the RNA interface (Figure 4F). This displacement exceeds the geometric constraints required for stable hydrogen bonding (< 3.5 Å), effectively abolishing its interaction with RNA. Across the interface, similar side-chain rearrangements (average displacement ∼3.07 Å, TableS2) propagate through the RNA-binding surface, reflecting transmission of allosteric stress from the drug-binding pocket.

As a direct consequence of these localized geometric perturbations, residue interaction network (RING) analysis ^51^ of the computational ensembles predicts a collapse of the hydrogen-bonding network between RNA and protein, with 24 hydrogen bonds lost upon drug binding (Figure 4G and TableS3). Key anchoring residues, including Lys8, Arg114, and Lys179, are reduced from multi-contact hubs to weak interactions, often shifted away from the iCLIP-defined RNA footprint. Several residues within RNA-XL-MS-identified RNA-binding regions (e.g., Asn14, Arg18, Arg307) completely lose RNA contacts, consistent with the disappearance of XL-MS signals at regions 9-41 and 298-311 (Figure 4G and 3G).

Together, these results demonstrate that duloxetine binding induces long-range allosteric remodeling of PyrB, leading to electrostatic repatterning and structural collapse of the RNA-binding interface, thereby disrupting RNA-protein interactions.

### RNA modulates the competitive inhibition of PyrB by duloxetine

To determine how the structural disruption induced by duloxetine translates into functional consequences, we performed enzyme kinetic assays to characterize PyrB activity. As mentioned above, PyrB catalyzes the rate-limiting step of *de novo* pyrimidine biosynthesis. Specifically, it catalyzes aspartate to N-carbamoyl-L-aspartate. Through targeted mass-spectrometry measurements, we monitored the concentration of N-carbamoyl-L-aspartate across varying concentrations of the substrate aspartate and the inhibitor duloxetine. We first performed preliminary kinetic profiling by fitting the data for each inhibitor concentration separately to the Hill equation (Figure 5A, main). This initial analysis revealed a pattern suggestive of mixed inhibition, with a dose-dependent increase in the half-saturation constant K_half_ (from 25.41 to 40.44 µM) and a concurrent decrease in the maximum velocity V_max_ (from 13.68 to 8.85 µM/min) (TableS4).

**Figure 5.**
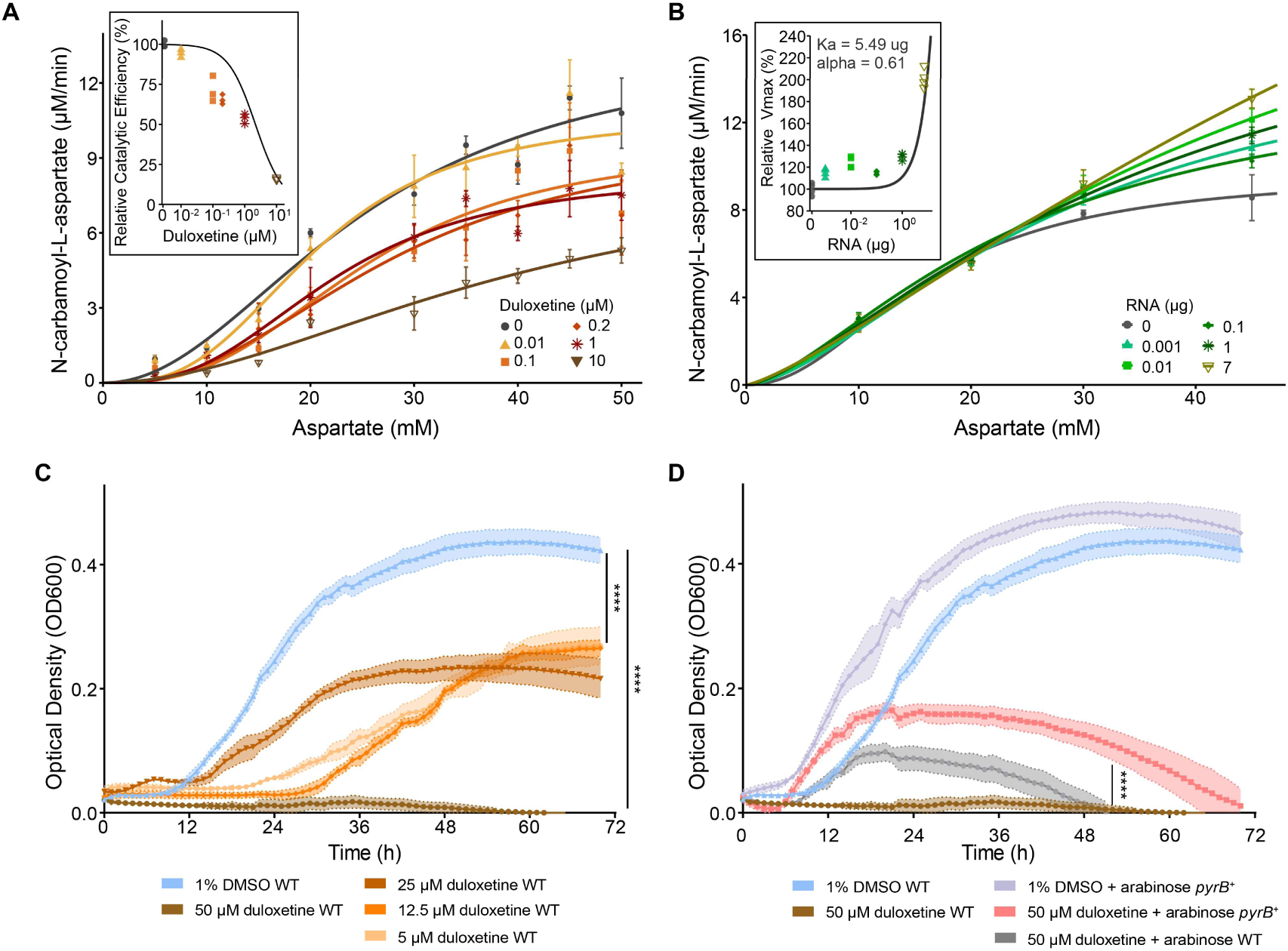
Duloxetine-binding inhibits the enzyme activity of PyrB, leading to the growth defect. **A.** Enzyme kinetic assays of recombinant *E. coli* IAI1 protein PyrB exposed to increasing concentrations of duloxetine (PyrB = 0.2 ng/mL). Individual Hill fits (lines) with mean ± SD of three biological replicates (points) are shown. Inset, relative catalytic efficiency (V_max_/K_half_, normalized to the no-inhibitor condition) as a function of duloxetine concentration, fitted with a competitive inhibition model (*K*_i_ = 2.16 µM). **B.** Enzyme kinetic assays of recombinant *E. coli* IAI1 protein PyrB exposed to increasing concentrations of total RNA with duloxetine held constant at 0.1 µM (PyrB = 0.2 ng/mL). Individual Hill fits (lines) with mean ± SD of four biological replicates (points) are shown. Inset, relative maximum velocity (V_max_, normalized to the RNA-free condition at 0.1 µM duloxetine) as a function of RNA concentration, fitted with the global mixed allosteric activation model. **C.** Growth curve of *E. coli* IAI1 on M9+1% glycerol media with different duloxetine concentrations (n = 6 biological replicates). The effect of growth inhibition was calculated using one-way ANOVA (F (4, 25) = 556.1, P < 0.0001, degrees of freedom 25, Dunnett’s t test). WT, wild type. **D.** Growth curve of *E. coli* IAI1 wild type (WT) and PyrB overexpression IAI1 strain (*pyrB*^+^, 0.01% arabinose as inducer) on M9+1% glycerol media (n = 6 biological replicates). The effect of growth inhibition was calculated using one-way ANOVA (F (4, 25) = 1024, P < 0.0001, Tukey’s test).

To resolve the precise mechanism and calculate a robust inhibition constant (K_i_), we performed a global analysis, fitting all data simultaneously to a cooperative competitive inhibition model (methods). This model yielded a good fit to the entire dataset (R² > 0.89) and confirmed significant positive cooperativity of PyrB, with a Hill coefficient (h) of 2.33 (95% CI, 1.83 to 2.94). Importantly, the analysis converged on a single intrinsic V_max_ of 10.85 (95% CI, 9.54 to 13.39) and a K_half_ of 23.81 µM (95% CI, 20.37 to 30.69). The relative catalytic efficiency declined dose-dependently with duloxetine concentration, yielding a K_i_ of 2.16 µM (95% CI, 1.11 to 3.93) (Figure 5A, inset; TableS4). These results strongly indicate that the primary mechanism of duloxetine inhibition is competitive, thereby compromising substrate binding.

Given that duloxetine and RNA appear to have competing effects on PyrB, we hypothesized that RNA could counteract the drug inhibition. To test this, we introduced a total RNA extract from *E. coli* IAI1 into the duloxetine-inhibited reaction (0.1 µM). Remarkably, the addition of RNA led to a potent, dose-dependent rescue of PyrB, with concurrent increases in both V_max_ and K_half_, a kinetic signature consistent with a mixed allosteric activation mechanism (Figure 5B and TableS4). To resolve the mechanistic basis of this RNA-mediated rescue, all kinetic data were simultaneously fitted to a mixed allosteric activation model with previously determined K_i_ at 2.16 µM (Methods). The global fit showed an RNA affinity constant K_a_ = 5.49 µg, and activation factor α = 0.61. At the highest RNA concentration tested (7 µg), the effective maximum velocity reached ∼200% of the inhibited baseline (Figure 5B, inset). We also found that RNA binding elevates V_max_ by a proportionally smaller factor than it raises K_half_ (TableS4), meaning that the enzyme achieves higher catalytic throughput at the cost of reduced substrate binding affinity. Together, these results indicate that RNA binding shifts PyrB into a distinct kinetic state with higher catalytic turnover but reduced substrate affinity, consistent with the previously observed conformational modulation in which RNA-binding creates an open catalytic state (Figure 2G and 4A).

Since PyrB catalyzes the rate-limiting step of *de novo* pyrimidine biosynthesis, inhibiting its activity should profoundly impact the growth of IAI1. To evaluate the *in vivo* efficacy of duloxetine, we conducted a growth assay in a pyrimidine-depleted defined medium (M9 + 1% glycerol), a condition under which PyrB function becomes essential for bacterial survival ^52^. Under the duloxetine concentrations ranging from 5 to 50 µM, a one-way analysis of variance (ANOVA) revealed a highly significant growth inhibition effect of the treatment on bacterial growth (F (4, 25) = 556.1, P < 0.0001). Specifically, under the physiologically relevant 50 µM duloxetine concentration ^6,12^, IAI1 growth almost completely ceased (Figure 5C). While overexpressing PyrB rescued the growth of IAI1 under a fully inhibitory duloxetine concentration of 50 µM (Tukey’s test, P < 0.0001, Figure 5D). This rescue was partial, however, as the growth of the *pyrB^+^* strain in the presence of the drug remained significantly lower than that of the uninhibited DMSO control (P < 0.0001). Together, these results suggested that PyrB is the functional target of duloxetine. The drug halts pyrimidine biosynthesis by directly competing with the natural substrate,

### Structural conservation of metabolic RBPs extends duloxetine’s off-target effect across diverse bacteria

PyrB is not only a critical metabolic enzyme and novel RBP in *E. coli*, but it is also essential to major bacterial pathogens, including *Helicobacter pylori* ^53^ and *Salmonella typhimurium* ^54^. To elucidate the structural conservation and taxonomic distribution of PyrB, we performed structural similarity searches using Foldseek ^55^. By searching against the manually curated AFDB-SWISSPROT database across bacteria and human, we confirmed the widespread structural conservation of high-confidence, annotated PyrB orthologs and their structural relatives (Ornithine Carbamoyltransferase, Figure S6A and TableS5) across diverse taxa, including the bioaccumulating *L. paracasei* we tested. Although *B. uniformis* and *S. salivarius* are absent from this database—a common limitation for non-model commensals—they both encode PyrB orthologs with highly conserved structural topologies predicted by AlphaFold (core fold root-mean-square deviation [RMSD] < 1 Å, Figure S6B).

Sequence-based phylogenetic analysis of these structurally similar proteins revealed strong conservation among gut-associated Enterobacteriaceae, including common pathogens such as *Salmonella*, *Shigella*, *Klebsiella*, and *Yersinia* (Figure 6A). Notably, when mapped onto this phylogeny, the human PyrB homolog (the aspartate transcarbamylase domain of the multifunctional protein CAD, UniProt ID: P27708) branched within the same clade as *E. coli*, highlighting an unexpected degree of similarity between the bacterial and host enzymes (Figure 6A and 6B). Crucially, human CAD is already an established RBP ^25,29,56^, with its RNA-interacting peptide aligning with the PyrB RNA-binding interface captured by our RNA-XL-MS data (AA85-106).

**Figure 6.**
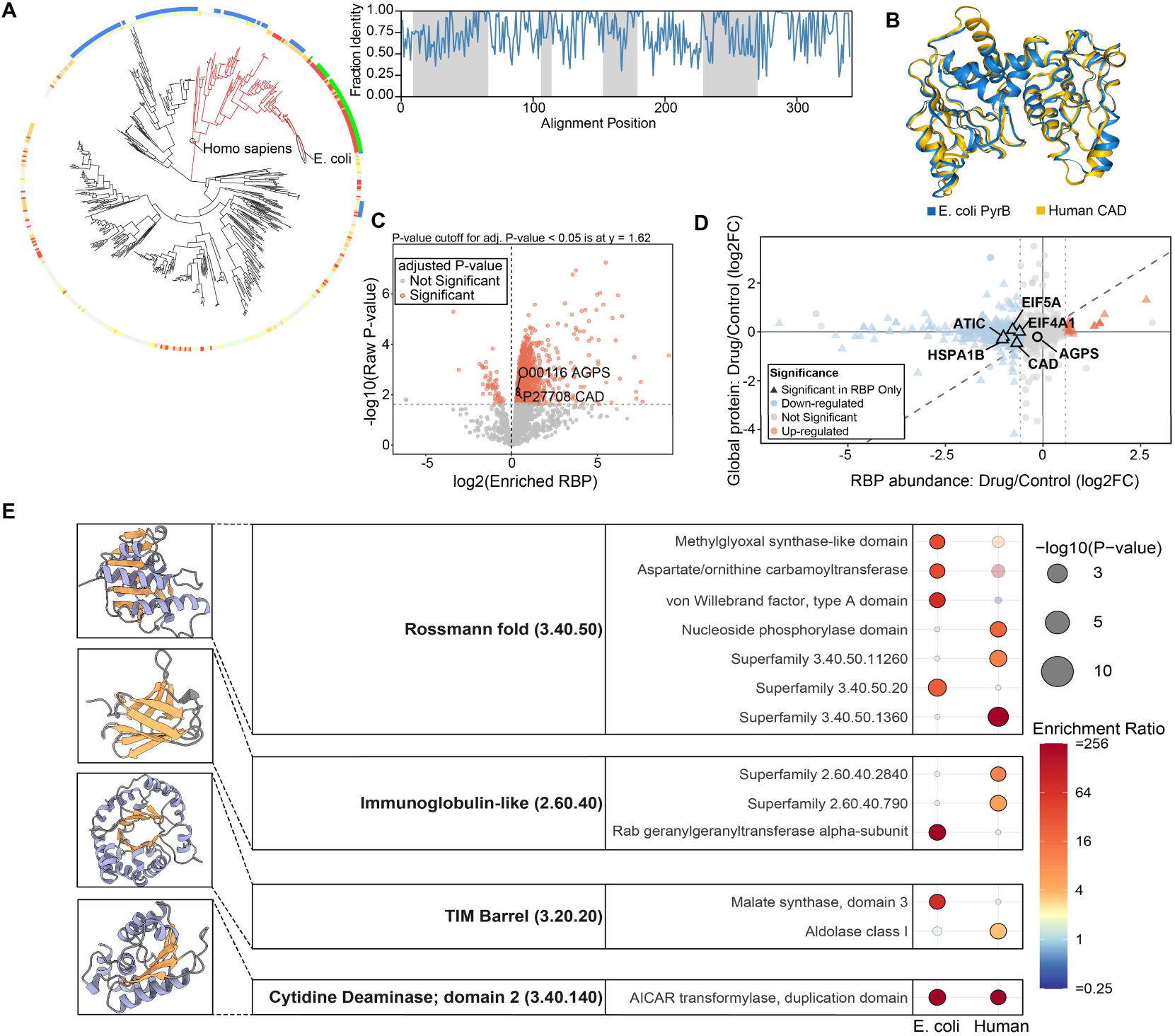
Structural conservation drives the pervasive cross-kingdom off-target effects of duloxetine. **A.** Phylogenetic distribution of PyrB across bacterial lineages. The outer ring color-coding denotes phylogenetic groups (green: *Enterobacteriaceae* species; blue: other gut-associated taxa). The inner track indicates the fraction of highly conserved residues (> 40% identity) within functionally critical peptide regions (more red means more conserved). The red-highlighted clade corresponds to the subset selected for sequence identity analysis (right inset graph). The inset graph quantifies the fraction identity across the alignment positions of these selected homologs, with grey shaded areas highlighting functionally important sequences that overlap with the multi-omics data in Figure 3F. **B.** The 3D structural alignment of *E. coli* PyrB (blue) and the functionally related domain of its human homolog, CAD (yellow). The overlay demonstrates high topological conservation, evidenced by a root-mean-square deviation (RMSD) of 1.53 Å and a Template Modeling (TM) score of 0.94. **C.** Identification of the RNA-binding proteome in human Caco-2 cells. The volcano plot highlights proteins significantly enriched in the RNase-negative fraction relative to the RNase-positive control (FDR-adjusted P < 0.05). **D.** Duloxetine selectively alters RNA-binding independent of global protein abundance. Scatter plot comparing the Log_2_ fold changes of the global protein abundance (y-axis) and the RNA-binding proteome (RBP abundance, x-axis) under duloxetine treatment compared to the control. The diagonal dashed line represents a 1:1 correlation of proportional change. The significantly up-regulated RNA-binding proteins are in red, while the significantly down-regulated RNA-binding proteins are in blue (FDR-adjusted P < 0.05, absolute log_2_ fold change in RBP abundance > 0.58). Triangles specifically designate proteins that exhibit significant alterations in RNA-binding without corresponding significant changes in their global protein expression levels (FDR-adjusted P > 0.05). **E.** CATH structural domain enrichment analysis. Comparison of duloxetine-disrupted RNA-binding proteins in *E. coli* IAI1 and human Caco-2 cells reveals shared structural vulnerabilities. Representative structural schematics for each domain are shown on the left, where α-helices are depicted in purple and β-sheets in beige. Node size corresponds to statistical significance (-log10 P-value), and node colour intensity indicates the enrichment ratio.

Within this shared clade (n=142 sequences), we observed an average of 72% sequence similarity, with 44 residues displaying 100% amino acid identity. These strictly conserved residues map directly to the drug- and RNA-interacting sites identified by our LiP-MS and RNA-XL-MS data (Figure 6A and 3G). This includes critical drug-binding anchor points (AA 53-56 STRT, R106, T169, R230, P269) and specific RNA-interacting sites (E87, P196). This conservation highlights a strong structural and evolutionary constraint on this specific RNA/substrate-binding domain.

Moreover, the disruptive effect of duloxetine is not limited to PyrB. Glycolate oxidase subunit GlcD, another prominent target from our OOPS-DIA dataset (Figure S2C), also exhibited duloxetine-disrupted RNA-binding, which we confirmed by EMSA (Figure S5A and S5B). Given that GlcD performs the vital function of oxidizing glycolate to glyoxylate, a process essential for *E. coli* survival on glycolate as a sole carbon source, we assessed the growth of IAI1 in M9-glycolate medium. Treatment with 50 µM duloxetine resulted in significant growth inhibition (one way ANOVA, F = 10.06, P < 0.0001, Dunnett’s t test), which can be rescued by GlcD overexpression, establishing it as a key physiological target of duloxetine during glycolate metabolism (Figure S5C and S5D). Collectively, these findings demonstrate that duloxetine exploits deep structural vulnerabilities shared by crucial metabolic RBPs like PyrB and GlcD. Given that these structural frameworks are highly conserved from gut bacteria to human homologs, we hypothesized that this drug-induced RBP disruption could transcend microbial systems and similarly impact the host.

### Duloxetine disrupts the RNA-binding proteome of human intestinal epithelial cells through cross-kingdom conserved structural domains

Bridging our investigation from microbial systems to the host, we confirmed that human intestinal epithelial cells (Caco-2) accumulate duloxetine (Figure S6C). Quantitative RNA-binding proteome revealed that specific human homologs, notably CAD (Figure 6B) and AGPS (a structural analog of *E. coli* GlcD protein) experienced impaired RNA associations by duloxetine (Figure 6C and S6D). Similar to *E. coli* IAI1, duloxetine triggered a global remodeling of the host cell RNA-binding proteome, significantly disrupting RNA-binding capacities of a distinct cohort of proteins without affecting their total protein abundances (Figure 6D and TableS6). Beyond generalized disruptions to structural cellular networks in epithelial barrier function and cytoskeletal organization (Figure S6E and S6F), duloxetine also disrupted the RNA-binding of three core translational regulators: HSPA1B, eIF4A1, and eIF5A (Figure 6D). These proteins represent sequential bottlenecks in the translation machinery, governing tRNA transcription ^57^, translation initiation ^58^, and elongation ^59^, respectively, suggesting that the host protein synthesis machinery could be systematically disrupted upon duloxetine treatment.

Given this widespread functional disruption across disparate pathways, we hypothesized that these proteins might share a common structural vulnerability. CATH domain enrichment analysis ^41^ of duloxetine-sensitive RBPs in both *E. coli* IAI1 and human Caco-2 cell revealed common structural targets, including the Rossmann fold (3.40.50), Immunoglobulin-like (2.60.40), TIM barrel (3.20.20), and cytidine deaminase; domain 2 (3.40.140). Notably, we identified the AICAR transformylase duplication domain—a core component of purine biosynthesis—as a conserved off-target in both bacterial (PurH) and human datasets (ATIC) (Figure 6E and TableS7). Structural analysis shows that this domain’s conserved deep hydrophobic cleft, which naturally binds AICAR and folate cofactors ^60^, perfectly accommodates the lipophilic naphthalene and thiophene rings of duloxetine (Figure S6G). We further characterized these enriched domains by mapping them to their primary biological functions and structural modes of action (Figure S7), which revealed that while these targeted proteins govern diverse processes—ranging from cytoskeletal organization to nucleotide metabolism—they share common structural vulnerabilities: hydrophobic pockets, protein interfaces, and lipophilic membrane anchors.

Taken together, these findings demonstrate that duloxetine-mediated disruption of RNA-protein interactions is not restricted to a single bacterial target but extends across conserved metabolic RBPs and diverse biological systems. Consistent with this broad distribution, duloxetine targets structurally conserved protein folds and disrupts RNA-binding regulatory networks present across a wide array of bacteria, including critical pathogens, as well as in human intestinal cells. These results reveal a shared molecular basis for drug-induced disruption of riboregulatory networks and highlight RNA-binding proteins as an overlooked layer of off-target drug activity.

## Discussion

Understanding how drugs impact biomolecular networks is essential for unraveling mechanisms underlying inter-individual variations in drug efficacy and toxicity. Duloxetine, widely used antidepressant, is a relevant case in point ^22,61–63^, warranting in-depth mechanistic investigation. Previous studies have largely focused on its classical protein targets involved in neurotransmission or metabolic regulation ^21,64,65^, whereas our findings bring forward protein-RNA interactions as a previously unrecognized layer of drug susceptibility. Rather than merely perturbing isolated targets, the scale of the observed disruption in these interactions is extensive. Across phylogenetically diverse bacteria and human intestinal cells, duloxetine consistently reduced RNA-binding capacity, ∼80% of RNA-binding proteins responded to duloxetine in bacteria (*E. coli* IAI1) and human (Caco-2) cells. At the cellular level, this high degree of response is greater than the dynamics of RNA-binding observed during major physiological transitions, such as cell cycle progression and growth phase shifts ^25,32^, or stress response ^29,66^, suggesting a deep impact on cellular molecular physiology. In our study, this impact is exemplified by the discovery of duloxetine-driven riboregulation of PyrB.

Aspartate transcarbamoylase (ATCase), with PyrB as its catalytic subunit, is a textbook model for multi-subunit allosteric regulation ^67,68^. Our study offers new insight by revealing a hidden layer of riboregulation that reshapes its kinetic response. We find that PyrB alone acts as a specific structural RNA sensor may coordinate its own substrate and energy networks. Its RNA targets, including transcripts encoding the TCA cycle enzyme SucD and the aspartate transporter DctA, suggest a novel regulatory circuit linking nucleotide biosynthesis with central carbon metabolism. This positions PyrB within the expanding landscape of moonlighting RNA-binding proteins, where metabolic enzymes orchestrate post-transcriptional networks ^23,24,27,31,35,69,70^. Our data shows that these riboregulatory hubs are vulnerable to xenobiotics: by simultaneously attenuating enzymatic activity and dismantling the riboregulatory complex, duloxetine acts via a new off-target mechanism distinct from classic active-site inhibition.

The implications of PyrB riboregulation extend across taxa due to its broad phylogenetic conservation. Comparing bacterial PyrB with the corresponding ATCase domain of its human ortholog (CAD), separated by billions of years of evolution, PyrB shows high structural conservation (RMSD = 1.53 Å). Beyond PyrB, we observe an enrichment of universal motifs like Rossmann folds and TIM barrels in the duloxetine-disrupted RNA-binding proteome. Such ancient folds are widespread among metabolic enzymes and RNA-binding proteins across both bacteria and human cells ^24,31,71–73^. These pervasive responses indicate that the structural vulnerabilities exploited by the drug are both broadly and deeply conserved.

As systemically circulating drugs extensively interact with the gut microenvironment, this widespread structural vulnerability offers a new molecular perspective on duloxetine’s common gastrointestinal side effects ^22^, and microbial disruptions ^74^. Given the established role of microbiome composition within the gut-brain axis in dictating antidepressant response ^75–78^, disruption of microbial riboregulatory networks raises an important hypothesis regarding the link between drug exposure, microbial state, and therapeutic variability. Moving forward, it will be critical to not only validate this conceptual framework through *in vivo* models, but also to evaluate other bioaccumulated molecules with similar physicochemical properties. Such investigations will clarify the broader prevalence of small-molecule mediated RBP modulation and unintended side effects.

Taken together, our study uncovers disruption of RNA-protein interactions as a hitherto hidden mechanism of off-target drug action. The currently known RBPs targeting molecules are designed for the purpose and therefore specific. Further, they compete at canonical RNA-binding interfaces or that engage the RNA itself ^79^. In contrast, our study reveals off-target and network-wide effect of an existing drug molecule. Biochemical and structural insights from our data suggest that such unselective drug-RBP disruptions are likely to be widespread and allosterically regulated. Because moonlighting RBPs couple metabolic and post-transcriptional control across kingdoms, allosterically accessible sites may represent a broad and largely unexploited druggable space. Moving forward, integrating structure-based computational screening with high-throughput RNA-interactome profiling and structural proteomics (such as OOPS-DIA and LiP-MS) could establish a powerful predictive pipeline. Incorporating the stability of these riboregulatory networks as a critical dimension in preclinical safety screening will be essential for developing microbiome-informed therapeutics with minimized off-target effects.

## Data availability

The mass spectrometry proteomics data have been deposited to the ProteomeXchange Consortium with the dataset identifiers PXD079294 (*E. coli* RNA-binding proteome, OOPS-DIA), PXD079520 (Human Caco-2 cell RNA-binding proteome, OOPS-DIA), PXD079475 (RNA-XL-MS), and PXD079876 (LiP-MS).

Targeted metabolomics raw data is deposited to Mendeley Data with DOI: 10.17632/x75bvvrbsb.1, accessible at https://data.mendeley.com/drafts/x75bvvrbsb.

iCLIP sequencing data have been deposited in the European Nucleotide Archive (ENA) under accession number PRJEB113415.

## Code availability

No custom algorithms were used for data analysis. The code used is available via GitHub at https://github.com/jiangxiaoteng/RBPome.

## Declaration of AI-assisted technologies used

During the preparation of this manuscript, the authors used ChatGPT and Gemini for spelling and grammar corrections. After using these tools, the authors reviewed and edited the content, and take full responsibility for the final manuscript.

## Author contributions

KRP, KSL and XJ conceptualized and designed the research. SM performed the phylogenetic analysis. IR, MM, SKP, and SK contributed to omics sample preparation. WZ contributed to sequencing data analysis. EV contributed to OOPS experiments. NB-C performed the cell culture experiments. AEL contributed to drug bioaccumulation analysis. KDI and NZ helped with recombinant protein expression. SB contributed to microbiology experiments. XJ wrote the first draft. XJ, KRP and KSL revised the manuscript. All authors read and commented on the final manuscript.

## Acknowledgements

We thank EMBL Genecore facility (Heidelberg, Germany) for iCLIP sequencing. This project has received funding from the European Research Council under the European Union’s Horizon 2020 research and innovation programme (grant no. 866028 to K.R.P), and from the UK Medical Research Council (project no. MC_UU_00025/11 to K.R.P.).

## Methods

### Gut bacteria anaerobic culture

Isolated gut bacteria *E. coli* IAI1, *E. coli* ED1a, *B. uniformis*, *L. paracasei*, and *S. salivarius* were grown as liquid cultures in mGAM at 37°C under anaerobic conditions (oxygen below 20 ppm, 15% carbon dioxide and 1.8–2% hydrogen). For duloxetine treatment, cell cultures from mGAM second passages were treated with 50 μM duloxetine or 0.1% DMSO control for 24 hours.

### Orthogonal Organic Phase Separation (OOPS)

A total of 2×10^7^ cells were collected for each replicate. RNA-binding proteins were extracted based on previously published ^25,32,34^, with minor adaptations. Cells were pelleted at 4,000×g for 15 min, then resuspended with 5 mL PBS and transferred to a 5 cm peri-dish in the anaerobic chamber. To covalently capture native RNA-protein complexes in situ, UV crosslinking was performed at 254 nm with 700 mJ/cm^2^ (CL-1000 Ultraviolet Crosslinker; UVP).

After crosslinking, cells were pelleted again to remove PBS and lysis by beating at 6000 rpm, 4×30 seconds on/off, 4 min, at 4 °C (MP FastPrep-24 5G Tissue Homogenizer; MP Biomedicals). 1 mL of acidic guanidinium-thiocyanate phenol (Trizol, Thermo Scientific) was added to each sample and vortexed for 15 seconds. Beads and cell debris were removed after centrifugation at 8,000×g 10 min at 4°C. The supernatant was transferred to a RNase-free tube and 100 μL of the total lysate was taken for the whole proteome analysis.

For the remaining cell lysate, 200 μL of chloroform (Fisher Scientific) and 1 ng of an external glycoprotein, human serotransferrin (UniProt ID: P02787), used as a quantification standard, were added to each *E. coli* sample. The mixture was vortexed for 10 seconds and then centrifuged at 12,000× g for 15 minutes at 4 °C. The aqueous phase containing non-crosslinked RNA was kept for further analysis. During this organic-aqueous phase separation, the crosslinked RNA-protein complexes were highly enriched at the interphase. The interphase containing RNA-protein complex and glycoprotein standard was subsequently washed two more rounds of Trizol phase separation, and then precipitated using 900 μL methanol. The pellete was washed two more times, air dried, and then resuspended using RNase-free water for RNase treatment.

At this stage, the completely dissolved interphase was divided into three fractions. One fraction was subjected to Proteinase K treatment to recover protein-bound RNA. The remaining two fractions were dedicated to quantitative proteomics to enable the identification of RNA-dependent proteins based on their selective depletion upon RNase digestion. The detailed steps for collecting non-crosslinked RNA and recovering protein-bound RNA were performed according to a previously published protocol ^34^.

For the two proteomics fractions, the RNase treatment was based on previously published ^32^, with minor modification. The interphase aliquots (90 μL each) were processed in parallel: one for RNase treatment, the other one as the RNase-untreated control. For the RNase-untreated sample, 10 μL 10× RNase control buffer (100 mM Tris-HCl pH 7.5, 3 M NaCl, 50 mM EDTA, 10× DTT) was added. For the RNase-treated sample, 10 μL 10X RNase digestion buffer (100 mM Tris-HCl pH 7.5, 3 M NaCl, 50 mM EDTA) and 2 μL of RNase A/T1 mix (Thermo Scientific, EN0551) were added. All samples were incubated for 4 h at 400 RPM and at 37 °C, followed by another around of the OOPS workflow. Interphases from both RNase-treated and RNase-untreated samples were pelleted and subjected to proteomics analysis.

### Proteomics sample preparation and LC-MS/MS analysis

The OOPS interphases were dissolved in 100 μL of 100 mM TEAB (Sigma-Aldrich) followed by DTT and IAA treatment and overnight trypsin digestion as previously reported ^34,80^. Samples were then acidified with trifluoroacetic acid (TFA, 0.1% [v/v] final concentration; Sigma-Aldrich) and the resulting peptides were subjected to C18 clean-up (C18 desalting columns, Thermo Scientific, 89851) following the manufacturer’s protocol.

The cleaned peptides were measured with bottom-up MS by using data-independent acquisition (DIA) with the following settings: The peptides were separated on a Dionex UltiMate 3000 system, and DIAMS acquisition was performed on an Orbitrap Eclipse Tribrid mass spectrometer (Thermo Fisher Scientific, San Jose, CA). The solvent system consisted of 0.1% FA in H_2_O (solvent A), and 80% ACN, 0.1% FA in H_2_O (solvent B). Proteolytic peptides (100 ng) were loaded onto an Acclaim PepMap 100 C18 trap column (0.1×20 mm, 5 µm particle size; Thermo Fisher Scientific) for 5 min at 5 µL/min with 100% solvent A. Peptides were eluted on an Acclaim PepMap 100 C18 analytical column (75 µm×50 cm, 3 µm particle size; Thermo Fisher Scientific) at 300 nL/min using the following gradient of solvent B: 2% for 5 min, linear from 2% to 20% in 125 min, linear from 20% to 32% in 40 min, up to 80% in 1 min, 80% for 9 min, down to 2% in 1 min, and 2% for 29 min, for a total gradient length of 210 min. Full MS spectra were collected at 120,000 resolution (AGC target: 3e6 ions, maximum injection time: 60 ms, 350-1,650 m/z), and MS2 spectra at 30,000 resolution (AGC target: 3e6 ions, maximum injection time: Auto, NCE: 27, fixed first mass 200 m/z). The isolation scheme consisted of 26 variable windows covering the 350-1,650 m/z range with an overlap of 1 m/z.

RAW data was processed using Spectronaut (version:17.0.200601.47784, Biognosys) with the following parameters: Prior to library-based analysis of the DIA data, the DIA raw files were converted into htrms files using the htrms converter. MS1 and MS2 data were centroided during conversion, and the other parameters were set to default. For the sample, the htrms files were analyzed with Spectronaut. Protein identifications were performed by searching against the respective UniProt reference proteomes: *E. coli* strain IAI1 (Proteome ID: UP000000746, download date: 2023_11_08), *E. coli* strain ED1a (Proteome ID: UP000000748, download date: 2023_12_12), and *Homo sapiens* (Proteome ID: UP000005640, download date: 2024_07_15). Quantitative analysis was performed by processing protein peak areas determined by the Spectronaut software. The identified common contaminant proteins were removed from the result lists before further analysis. Normalization was based on the added serotransferrin followed by DEqMS for differential protein expression in quantitative proteomic analysis ^81^. The significantly enriched proteins in RNase-untreated samples are defined as RNA-binding proteins, since they are sensitive to RNase treatment in OOPS ^34^.

### Recombinant protein expression and purification

The genes encoding the proteins of interest, *pyrB* and *glcD*, were amplified and cloned into the expression vector pBADHis_A (Thermo Scientific) to enable arabinose-inducible regulation of protein expression. The recombinant vector introduces a cleavable N-terminal polyhistidine tag, which was then transformed into E. coli BL21 and E. coli IAI1 for protein purification and iCLIP experiment, respectively.

Protein expression and purification was based on the standard procedure ^82^, with different levels of L-arabinose induction to achieve an optimal result. For the purification of PyrB, final concentration of 0.02% arabinose was applied to the LB culture and induced at 30°C for 4 hours. The bacteria were pelleted and lysed by sonication (50% intensity, 10s on/20s off, 5 min total run time) in lysis buffer (50 mM Tris-HCl pH 7.4, 150 mM NaCl, 5% glycerol, 1 mM DTT, 40 mM imidazole, 1× cOmplete protease inhibitor cocktail, 10U RNase A, and 10U Turbo DNase). HisTrap HP column on an AKTA go protein purification system was used to purify the his-tagged protein. The purified protein was verified using stain-free gels and western blot.

### Electromobility shift assay (EMSA)

Total RNA from *E. coli* IAI1 was extracted using the PureLink RNA Mini Kit (Thermo Scientific) according to the manufacturer’s instructions, to assess the binding of PyrB and GlcD, respectively. The binding reaction was carried out in a 20 μL mixture containing protein (0.4-12 μg, protein gradient assay), RNA (1-17 μg, RNA gradient assay), and duloxetine (0-6000 μM, drug gradient assay), in a binding buffer (4 U/μL RNAsin, 5 mM DTT, 0.5 mM PMSF, 2.5 mM MgCl_2_, 100 mM KCl; 20 mM HEPES, pH 7.5; 0.2 mM EDTA and 20% glycerol) modified from previously published protocols ^27^. The reactions were incubated at 25°C with shaking at 300 rpm for 20 min on a thermomixer. Then the reaction was mixed with 4 μL 6X native loading buffer and loaded on the native gel (4%-15% polyacrylamide gel, 60 V, 2 hours). The RNA and proteins are stained sequentially in the gel after electrophoresis according to the manufacturer’s instructions (Thermo Scientific, E33075).

For the EMSA-Western Blot assay, the 20 μL reaction mixture is basically the same with less protein (0.3-0.6 μg) and RNA (2 μg or 8 μg) input. After gel electrophoresis, proteins were transferred to nitrocellulose membranes using the Trans-Blot Turbo Transfer System (Bio-rad) and followed the standard procedure. The nitrocellulose membrane was blocked with 3% milk in TBST for 1 hour at room temperature. Primary antibodies (1:5000) were incubated in 3% milk in TBST either overnight at 4°C or one hour at room temperature (RT), followed by 6 times TBST washes, secondary antibody (1:10000) incubation in 3% milk TBST for 1 hour at room temperature. Then the membrane was washed 6 times in TBST and developed using the ECL solution. The western blot signal was detected with the ChemiDoc System and analyzed with the Image Lab (Bio-Rad, version 6.1).

### Enzymatic activity assays

Targeted metabolomics was performed to assess the enzymatic activity of PyrB. The reaction mixture was constructed based on the previous paper ^49^, with minor adaptations. Briefly, the substrates L-aspartate and carbamoyl phosphate (Sigma) were dissolved in 50 mM Tris-Acetate. The reaction mix contained 0.2 ng/mL PyrB, L-aspartate at a final concentration of 0-50 mM, and duloxetine at a final concentration of 0-10 μM (or 0.1% DMSO as control). For the RNA gradient experiment, total RNA extract (0–70 μg/mL) was added instead of duloxetine. The mixture was equilibrated to 25 °C in a water bath for 10 min. followed by the addition of 2 μL 240 mM carbamoyl phosphate (final concentration 48 mM) to initiate the reaction in a total volume of 100 μL. The reaction was incubated at 37 °C for 30 minutes and then quenched by adding 200 μL of ice-cold metabolomics extraction buffer (1:1:1 acetonitrile:methanol:ddH2O with 75µM caffeine). After incubation at 4 °C for 10 minutes, the mixture was centrifuged at 4,000 × g for 3 min at 4 °C. The supernatant was collected for LC-MS/MS analysis.

Targeted metabolite quantification on N-carbamoyl-L-aspartate was performed using an Agilent 1290 Infinity II UPLC system coupled to an Agilent 6470 triple quadrupole (QQQ) mass spectrometer equipped with a JetStream electrospray ionization (AJS-ESI) source. Metabolite separation was achieved by hydrophilic interaction liquid chromatography (HILIC) using an Atlantis™ Premier BEH Z-HILIC column (1.7 µm, 2.1 × 100 mm, Waters) maintained at 40 °C. The injection volume was 0.3 µL, and the flow rate was set to 0.9 mL/min.

The mobile phases consisted of Buffer A (water with 10 mM ammonium bicarbonate) and Buffer B (90% acetonitrile with 10 mM ammonium bicarbonate). The elution gradient was as follows: 0–0.7 min, 85% B; 0.7–2.55 min, linear decrease to 50% B; 2.55–2.9 min, hold at 50% B; 2.9–2.91 min, return to 85% B; 2.91–3.2 min, re-equilibration. The total run time was 3.2 minutes.

Data was acquired in dynamic (scheduled) multiple reaction monitoring (MRM) mode with a cycle time of 320 ms. Source parameters were set as follows: gas temperature, 325 °C; gas flow, 10 L/min; nebulizer pressure, 40 psi; sheath gas temperature, 350 °C; sheath gas flow, 11 L/min; capillary voltage, ±3500 V; nozzle voltage, ±1000 V. Data acquisition was performed using MassHunter Workstation LC/MS Data Acquisition software (v10.1), and peak integration and quantification were conducted in MassHunter Quantitative Analysis for QQQ (v10.1).

The resulting quantitative data was fitted with the Hill equation as previously reported ^83^. Vmax, Khalf, and h were calculated from the Hill equation, and graphs were plotted by GraphPad Prism 8. The cooperative competitive inhibition model (allosteric sigmoidal kinetics with competitive inhibition) was generated using the following equation to resolve the precise inhibition mechanism and kinetic constants:

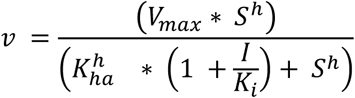

where the *v* stands for the initial velocity, *S* is the substrate concentration, *I* is the inhibitor concentration, V_max_ is the maximum velocity, K_half_ is the substrate concentration at half-maximal velocity, h is the Hill coefficient, and K_i_ is the inhibition constant.

The mixed activation model is defined by the equation:

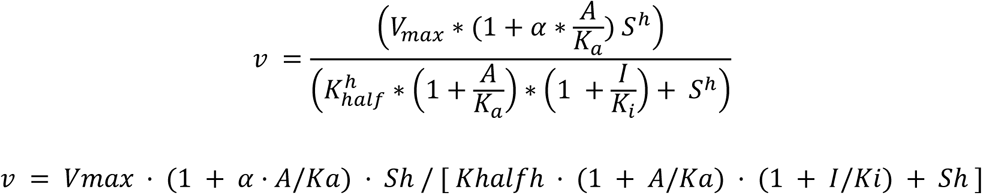

where *A* is the RNA concentration, *I* is the fixed duloxetine concentration (0.1 µM), and α is the activation asymmetry factor.

### Conditional growth assay

*E. coli* IAI1 wild-type (WT) and PyrB/glcD overexpression strains (PyrB⁺/glcD⁺) were cultured overnight in mGAM medium and subsequently passaged into M9 medium supplemented with 0.4% glucose for a second passage (P2). P2 cells were washed with PBS and inoculated into 96-well plates at an initial OD₆₀₀ of 0.05 in defined media (M9 supplemented with 0.4% glucose, 1% glycerol, or 120 mM glycolate), containing duloxetine at concentrations ranging from 5 to 50 μM (0.1% DMSO as control). For the overexpression strains, 0.01% arabinose was added to induce gene expression. Cultures were grown under anaerobic conditions at 37 °C for up to 48 hours, with OD₆₀₀ measurements taken hourly to monitor bacterial growth over time.

To quantify growth differences across conditions, the area under the curve (AUC) was calculated in GraphPad Prism 9 for each replicate. Statistical comparisons were performed using unpaired two-tailed Student’s t-tests, with biological replicates (n = 3–5) as indicated. All plots and statistical analyses were generated using GraphPad Prism 8.

### iCLIP

The iCLIP experiment was conducted based on the previously described protocol ^84^. *E. coli* IAI1 wild-type (WT) and PyrB overexpression strains (*pyrB⁺*) were grown in mGAM with 50 μM duloxetine or 0.1% DMSO under anaerobic conditions at 37 °C and induced with 0.01% L-arabinose at the exponential phase (OD ∼0.4) and harvested at stationary phase (OD ∼1.2). Then the cells were washed with PBS and UV crosslinked as described above. The UV crosslinked cells were resuspended in iCLIP lysis buffer (50 mM Tris-HCl pH 7.4, 100 mM NaCl, 1% Igepal CA-630 (Sigma I8896), 0.1% SDS, 0.5% sodium deoxycholate, 1× cOmplete™ protease inhibitor cocktail) and lysed using sonication (50% intensity, 10s on/20s off, 5 min total run time). Then the cell lysates were treated with 1 μL diluted RNase I (0.1-4 U/mL final concentration, EN0602, Thermo Scientific) to determine the optimal RNase treatment and with 2 μL TURBO DNase (AM2238 Thermo Fisher Scientific) to remove the unwanted DNA. RNA-protein complex was captured using HisPur™ Ni-NTA Magnetic Beads following the manufacturer’s instructions. RNA dephosphorylation, pre-adenylated L3-1R-App 3’ adaptors ligation and excess adaptors removal were carried out exactly as described ^84^. RNA-protein complexes were isolated by SDS-PAGE and nitrocellulose transfer. Pre-adenylated RNA signal was captured using the ChemiDoc System at both IRDye 680RD and 800CW channels. RNA was then released from the membrane by Proteinase K treatment and purified by phenol-chloroform extraction. Isolated RNA was reverse transcribed with SuperScript IV (Invitrogen, 18090010) and the resulting DNA was circularized and amplified by PCR. The cDNA library after PCR amplification (Phusion HF Master mix, New England Biolabs) was subjected to the purification with AMPure XP Beads (Beckman Coulter Life Sciences) following the manufacturer’s instructions. Then the purified cDNA library was subjected to Qubit quantification and Tape station analysis before sequencing on the Illumina NextSeq 2000 P2 100 bp in single-end mode. Adaptor and primer sequences are listed in TableS8. Sequencing data have been deposited in the European Nucleotide Archive (ENA) under accession number PRJEB113415.

iCLIP data analysis was performed based on the previously published paper ^85^ with minor adjustments based on the data format. Briefly, the UMI-extracted data was aligned to tRNA and rRNA sequences to filter out contaminations. The cleaned reads were mapped to the *E. coli* IAI1 genome GCF_000026265.1 using STAR with default parameters according to the reference ^85^. PCR duplicates were removed based on Unique Molecular Identifiers (UMIs) and mapping coordinates. Uniquely mapped reads were retained, and cross-link sites were defined as the nucleotide preceding the read start (start - 1). Peak calling was performed on the merged replicate samples using PureCLIP after cleaning sequencing reads less than 20 nt to remove low-confidence fragments and sequence artifacts. The downstream peak analysis was done using the BindingSiteFinder package ^85^. Peak visualization was generated using IGV ^86^. RNA binding motifs were detected based on Positionally enriched k-mer analysis (PEKA) ^87^ and MEME ^88^ with default parameters. The candidate RNA sequence secondary structures were predicted using RNAfold from the ViennaRNA (Version 2.7.2) centroid structure ^89^.

### Identification of RNA-binding sites

Crosslinking sites on our target protein PyrB were identified based on the NuXL method with some modifications ^50^. Briefly, for the in vitro binding test, the binding reaction was carried out in a 20 μL mixture containing 5 μg protein, 5 μg RNA, and duloxetine (final concentration 5000 μM), in a binding buffer same as EMSA assay above (4 U/μL RNAsin, 5 mM DTT, 0.5 mM PMSF, 2.5 mM MgCl_2_, 100 mM KCl; 20 mM HEPES, pH 7.5; 0.2 mM EDTA and 20% glycerol). UV crosslinking was performed the same as above. The crosslinked samples were precipitated using methanol and washed once with ice-cold 80% [v/v] ethanol. The pellets were resolved in 8 M urea in 50 mM Tris-HCl (pH 7.5) then diluted to 1 M urea using 50 mM Tris-HCl. The dissolved samples were treated with 250 U Pierce™ Universal nuclease for Cell Lysis (Thermo Scientific™, 88700) and 100 U nuclease P1 (NEB, M0660S), followed by incubation for 3 hours at 37°C, 300 rpm. Trypsin was added at a 1:25 enzyme-to-protein ratio, incubated overnight at 37°C, 300 rpm. Samples were acidified with formic acid to pH 5 then incubated with 100 U nuclease P1 for 30 min at 37°C, 300 rpm. Samples were acidified with formic acid to pH 3-4 followed by C18 clean-up as described above. The crosslinked peptide(oligo)nucleotides were enriched using TiO_2_ immobilized resins in spin column (Thermo Scientific™, A32993) as per manufacturer’s instructions and then subjected to LC-MS/MS measurements.

Data-dependent acquisition (DDA) was performed on an Orbitrap Exploris 480 Mass Spectrometer using a 60-minute gradient. The liquid chromatography system utilized 0.1% FA water solution as solvent A and 80% acetonitrile as solvent B. Peptides were loaded onto a trap column and separated on a 25 cm analytical column maintained at 40°C. Peptides were eluted at a flow rate of 300 nL/min using a 40-minute linear gradient from 2% to 40% solvent B, followed by a 5-minute linear gradient from 40% to 90% B and a 15-minute gradient from 90% to 95% B. Full MS spectra were acquired in the Orbitrap from m/z 370–1500 at a resolution of 120,000. The automatic gain control (AGC) target was set to a custom normalized value of 300% with a maximum injection time of 50 ms.

For MS2 analysis, the most intense precursor ions with charge states from 2 to 5 were selected for fragmentation. Precursors were fragmented using Higher-energy Collisional Dissociation (HCD) with a normalized collision energy (NCE) of 26%. The resulting MS2 spectra were acquired at a resolution of 15,000. The MS2 scan parameters included a custom normalized AGC target of 100%, a maximum injection time of 45 ms, and an isolation window of 2.0 m/z. The method operated with a total cycle time of 1.5 seconds, and dynamic exclusion was enabled for 65 seconds to prevent re-sequencing of the same precursor.

For in vivo crosslink and immunoprecipitation of target RNA-protein complex, *E. coli* IAI1 wild-type (WT) and PyrB overexpression strains (*pyrB⁺*) were grown in mGAM with 50 μM duloxetine or 0.1% DMSO under anaerobic conditions as described above. Then the target complex from UV crosslinked cells was enriched through HisPur™ Ni-NTA magnetic beads (Thermo Scientific™, 88831) as per manufacturer’s instructions. The enriched complex was subjected to precipitation, RNase digestion, tryptic digestion and TiO_2_ enrichment same as the above mentioned NuXL method.

### Limited proteolysis and LC-MS/MS (LiP-MS)

The LiP-MS experiment was conducted based on previously described ^48^ with a few adaptations. Briefly, 10 μg of purified PyrB protein was incubated in a 50 μL reaction mixture containing 20 mM HEPES (pH 7.5), 150 mM KCl, 10 mM MgCl₂, and 5% glycerol. Duloxetine (final concentration 5000 μM) or 0.1% DMSO (control) was added, and the mixture was incubated at 25 °C for 10 min. Subsequently, 5 μL of diluted proteinase K (1:100 w/w, enzyme-to-substrate ratio) was added to the LiP samples (LiP-drug and LiP-DMSO), while 5 μL of ddH₂O was added to the control samples (TrP-drug and TrP-DMSO). All samples were then incubated at 25 °C for 5 minutes, followed by inactivation of proteinase K at 99 °C for 5 minutes. Sodium deoxycholate (DOC) was added to a final concentration of 5% (w/v). Both LiP and TrP samples were subjected to complete tryptic digestion under denaturing conditions as described previously ^48^. After tryptic digestion, peptides were cleaned using detergent removal columns (Thermo Scientific, 87777) and C18 desalting columns (Thermo Scientific, 89851). To reduce false discovery rates (FDR) and prevent misannotation during database searching, a heterologous whole-cell HeLa digest was spiked into each sample at a 1:2 ratio (sample:HeLa digest).

Data-dependent acquisition (DDA) was performed to generate the spectral library on an Orbitrap Eclipse Tribrid mass spectrometer in the DDA mode. The LC method was applied using a total run time of 180 min gradient. The gradient of solvent B (80% ACN, 0.1%FA) was as follows: an initial condition of 2% B for 3 min, a linear gradient from 5% to 40% B in 142 min, linear from 40% to 45% B in 10 min, followed by a linear gradient from 45% to 90% B in 5 min. The gradient was held at 90% B for 5 min, then returned to 3% B and held at 3% B for the remainder of the run for column re-equilibration.

Full MS spectra were acquired in the Orbitrap from m/z 400–1600 at a resolution of 120,000. The Automatic Gain Control (AGC) target was set to Standard with the maximum injection time on Auto. Precursor ions with charge states from 2 to 6 were selected for fragmentation. Dynamic exclusion was active for 60 seconds after a single acquisition. Data-dependent MS2 spectra were generated using Higher-energy Collisional Dissociation (HCD) with stepped normalized collision energies of 25% and 30%. MS2 scans were acquired in the Orbitrap at a resolution of 50,000 with a fixed first mass of m/z 120. For MS2 scans, a custom 250% normalized AGC target and a 200 ms maximum injection time were utilized with an isolation window of 0.7 m/z.

Spectral library generation was performed based on previously reported ^48^ using Spectronaut (Biognosys) hybrid library generation features, with the default setting and matching the tryptic digestion rule of the search settings. DIA was performed for LFQ quantification using the same parameters as described above in OOPS proteomics data. Data processing was performed by using the R-base package MSstatsLiP ^48^.

### Molecular modeling of duloxetine-protein-RNA interaction

Protein structures were obtained from the Protein Data Bank (PDB) ^46,90^ and AlphaFold ^91,92^. Structural alignments were conducted in UCSF ChimeraX, with the Root Mean Square Deviation (RMSD) values calculated using default parameters ^93^. Molecular docking was performed to evaluate the interaction and binding affinity between target proteins and duloxetine following the default steps ^94^. Briefly, Small-molecule ligands (duloxetine) were prepared in AutoDockTools (v1.5.7) by adding hydrogen and defining torsion flexibility. Protein structures were preprocessed by removing solvent molecules and adding Gasteiger charges. Docking was carried out using AutoDock Vina with an exhaustiveness value of 28. In cases where the binding site was unknown, the docking grid was set to cover the entire protein surface (blind docking). Protein-ligand interactions, including molecular contacts and hydrogen bonds, were analyzed using ChimeraX 1.9 with default parameters.

For predicting the RNA-protein complex, the initial models were generated using AlphaFold and RoseTTAFold2NA ^95^. To account for RNA flexibility, the secondary structure was predicted via RNAfold ^89^ and used as structural constraints for RNAcomposer ^96^ and SimRNA ^97^ to construct a diverse 3D RNA conformational ensemble (n=4). Experimental XL-MS and iCLIP data, alongside 2D base-pairing information, were translated into ambiguous interaction restraints (AIRs) to drive the docking process.

Macromolecular docking was performed using HADDOCK 3.0 ^98^. For the RNA-protein model, the rigid-body docking phase underwent extensive sampling (10,000 iterations) to ensure adequate conformational exploration. This was followed by semi-flexible simulated annealing (flexref) and energy minimization refinement (emref) to generate reliable conformations based on the recommended settings ^98^. The generated complexes were clustered and evaluated using the HADDOCK scoring function. Because the molecular dynamics refinement performs a short molecular dynamics simulation and its computationally demanding, we chose the top 2 scored models from the top 3 cluster (a total of 6 representative models) for this step. The final generated models were subjected to an explicit-solvent refinement (water) phase, with a total 20 ps of molecular dynamics simulation for 10,000 integration steps at 300 K to relax the steric clashes and optimize the water-mediated interactions at the RNA-protein interface.

For the duloxetine-bound state, the explicit ligand topology and parameter files in CNS format were generated using the ACPYPE web server and integrated into the HADDOCK library ^99^. The protein, RNA ensemble, and the ligand duloxetine were input as three independent entities. The pre-docked protein-ligand complex containing protein-duloxetine interactions was extracted and set as specific distance restraints for the three-body docking. Other parameters were the same as above.

Interfacial binding free energies (ΔiG) were calculated using PDBePISA ^100^, while the buried surface areas (BSA) and geometric properties were quantified using ChimeraX 1.9 with default parameters. To map the allosteric disruption, non-covalent residue interaction networks (hydrogen bonds, van der Waals, etc.) were computed via the RING 3.0 server ^51^ and visualized using Cytoscape ^101^.

### Microfluidic Diffusional Sizing (MDS) for RNA-protein interactions

In-solution RNA-protein binding affinities and subsequent drug-induced dissociation were quantified using Microfluidic Diffusional Sizing (MDS) on a Fluidity One-M system (Fluidic Sciences) following the manufacturer’s guide. The system measures the hydrodynamic radius (R_h_) of fluorescent species based on its diffusion profile under steady-state laminar flow. Complex formation between the labeled probe and the unlabeled target results in a measurable increase in R_h_, enabling the calculation of the equilibrium dissociation constant (K_D_) using standard binding isotherms ^102^. All experiments were performed in RNase-free PBS buffer supplemented with 0.005% Tween-20 to prevent non-specific surface adsorption of the fluorescent tagged RNA.

For the RNA-protein binding assay, fluorescently labeled RNA probes were dissolved in nuclease-free TE buffer and diluted to a final working concentration of 20 nM in the assay mixture. Recombinant PyrB protein was prepared as a serial dilution series covering a concentration range sufficient to capture the full binding curve. The fluorescent RNA was mixed with the PyrB protein gradient and incubated 30 mins to reach binding equilibrium before MDS analysis.

To quantify the allosteric disruption of the RNA-binding interface by duloxetine, drug perturbation MDS assays were conducted under two scenarios. In the pre-inhibition scenario, a serially diluted gradient of duloxetine was incubated with PyrB first for 30 minutes before adding RNA for another 30 minutes incubation. In the dissociation scenario, the RNA-protein complexes were pre-assembled and subsequently treated with the duloxetine gradient. To eliminate solvent-induced artifacts, the final concentration of the drug vehicle (DMSO) was strictly maintained at 1% across all gradient points and the respective vehicle controls. Bovine serum albumin (BSA) was used as a negative control to verify interaction specificity. Binding data and complex fractions were fitted using the Levenberg-Marquardt nonlinear least-squares algorithm in R.

### Phylogenetic tree generation and residues conservation

The protein sequences of the Aspartate carbamoyltransferase that were selected based on structure conservation were aligned using MEGA (v12) and Muscle with default options. The alignment was visualised and inspected using Jalview (v2.11). Residues conserved in more than 40% of the genomes were marked for display in the phylogenetic tree. The phylogenetic tree was generated using the Jones-Taylor-Thornton model and visualised with iTOL ^103^.

## Reference

1. Aasmets, O. et al. A hidden confounder for microbiome studies: medications used years before sample collection. mSystems 10, e00541–25 (2025).

2. Falony, G. et al. Population-level analysis of gut microbiome variation. Science 352, 560–564 (2016).

3. Lindell, A. E., Zimmermann-Kogadeeva, M. & Patil, K. R. Multimodal interactions of drugs, natural compounds and pollutants with the gut microbiota. Nat. Rev. Microbiol. 20, 431–443 (2022).

4. Vieira-Silva, S. et al. Statin therapy is associated with lower prevalence of gut microbiota dysbiosis. Nature 581, 310–315 (2020).

5. Garcia-Santamarina, S. et al. Emergence of community behaviors in the gut microbiota upon drug treatment. Cell 187, 6346–6357.e20 (2024).

6. Maier, L. et al. Extensive impact of non-antibiotic drugs on human gut bacteria. Nature 555, 623–628 (2018).

7. Kumar, A. et al. Identification of medication–microbiome interactions that affect gut infection. Nature 644, 506–515 (2025).

8. Trepka, K. R., Olson, C. A., Upadhyay, V., Zhang, C. & Turnbaugh, P. J. Pharma[e]cology: How the Gut Microbiome Contributes to Variations in Drug Response. Annu. Rev. Pharmacol. Toxicol. 65, 355–373 (2025).

9. Verdegaal, A. A. & Goodman, A. L. Integrating the gut microbiome and pharmacology. Sci. Transl. Med. 16, eadg8357 (2024).

10. Balaich, J. et al. The human microbiome encodes resistance to the antidiabetic drug acarbose. Nature 600, 110–115 (2021).

11. Zimmermann, M., Zimmermann-Kogadeeva, M., Wegmann, R. & Goodman, A. L. Mapping human microbiome drug metabolism by gut bacteria and their genes. Nature 570, 462–467 (2019).

12. Klünemann, M. et al. Bioaccumulation of therapeutic drugs by human gut bacteria. Nature 597, 533–538 (2021).

13. Burke, K. & Li, Y. Impact of Gut Microbiota on Drug Metabolism and Absorption. Curr. Pharmacol. Rep. 11, 49 (2025).

14. Weersma, R. K., Zhernakova, A. & Fu, J. Interaction between drugs and the gut microbiome. Gut 69, 1510 (2020).

15. Zimmermann, M., Patil, K. R., Typas, A. & Maier, L. Towards a mechanistic understanding of reciprocal drug–microbiome interactions. Mol. Syst. Biol. 17, e10116 (2021).

16. Haiser, H. J. et al. Predicting and Manipulating Cardiac Drug Inactivation by the Human Gut Bacterium Eggerthella lenta. Science 341, 295–298 (2013).

17. Maini Rekdal, V., Bess, E. N., Bisanz, J. E., Turnbaugh, P. J. & Balskus, E. P. Discovery and inhibition of an interspecies gut bacterial pathway for Levodopa metabolism. Science 364, eaau6323 (2019).

18. Daniel, W. A. Mechanisms of cellular distribution of psychotropic drugs. Significance for drug action and interactions. Prog. Neuropsychopharmacol. Biol. Psychiatry 27, 65–73 (2003).

19. Kazmi, F. et al. Lysosomal Sequestration (Trapping) of Lipophilic Amine (Cationic Amphiphilic) Drugs in Immortalized Human Hepatocytes (Fa2N-4 Cells). Drug Metab. Dispos. 41, 897–905 (2013).

20. Tummino, T. A. et al. Drug-induced phospholipidosis confounds drug repurposing for SARS-CoV-2. Science 373, 541–547 (2021).

21. Abdi, S. A. H., Azhar, G., Zhang, X. & Wei, J. Y. Duloxetine, an SNRI, Targets pSTAT3 Signaling: In-Silico, RNA-Seq and In-Vitro Evidence for a Pleiotropic Mechanism of Pain Relief. Int. J. Mol. Sci. 26, 10432 (2025).

22. Knadler, M. P., Lobo, E., Chappell, J. & Bergstrom, R. Duloxetine: clinical pharmacokinetics and drug interactions. Clin. Pharmacokinet. 50, 281–294 (2011).

23. Castello, A., Hentze, M. W. & Preiss, T. Metabolic Enzymes Enjoying New Partnerships as RNA-Binding Proteins. Trends Endocrinol. Metab. 26, 746–757 (2015).

24. Hentze, M. W., Sommerkamp, P., Ravi, V. & Gebauer, F. Rethinking RNA-binding proteins: Riboregulation challenges prevailing views. Cell 188, 4811–4827 (2025).

25. Queiroz, R. M. L. et al. Comprehensive identification of RNA–protein interactions in any organism using orthogonal organic phase separation (OOPS). Nat. Biotechnol. 37, 169–178 (2019).

26. Hentze, M. W. & Preiss, T. The REM phase of gene regulation. Trends Biochem. Sci. 35, 423–426 (2010).

27. Huppertz, I. et al. Riboregulation of Enolase 1 activity controls glycolysis and embryonic stem cell differentiation. Mol. Cell 82, 2666–2680.e11 (2022).

28. Spizzichino, S. et al. Structure-based mechanism of riboregulation of the metabolic enzyme SHMT1. Mol. Cell 84, 2682–2697.e6 (2024).

29. Trendel, J. et al. The Human RNA-Binding Proteome and Its Dynamics during Translational Arrest. Cell 176, 391–403.e19 (2019).

30. Christopoulou, N. & Granneman, S. The role of RNA-binding proteins in mediating adaptive responses in Gram-positive bacteria. FEBS J. 289, 1746–1764 (2022).

31. Hentze, M. W., Castello, A., Schwarzl, T. & Preiss, T. A brave new world of RNA-binding proteins. Nat. Rev. Mol. Cell Biol. 19, 327–341 (2018).

32. Monti, M. et al. Interrogation of RNA-protein interaction dynamics in bacterial growth. Mol. Syst. Biol. 20, 573–589 (2024).

33. Rüttiger, A.-S. et al. The global RNA-binding protein RbpB is a regulator of polysaccharide utilization in Bacteroides thetaiotaomicron. Nat. Commun. 16, 208 (2025).

34. Villanueva, E. et al. Efficient recovery of the RNA-bound proteome and protein-bound transcriptome using phase separation (OOPS). Nat. Protoc. 15, 2568–2588 (2020).

35. Holmqvist, E. & Vogel, J. RNA-binding proteins in bacteria. Nat. Rev. Microbiol. 16, 601–615 (2018).

36. Hör, J. et al. Grad-seq shines light on unrecognized RNA and protein complexes in the model bacterium Escherichia coli. Nucleic Acids Res. 48, 9301–9319 (2020).

37. Shchepachev, V. et al. Defining the RNA interactome by total RNA-associated protein purification. Mol. Syst. Biol. 15, e8689 (2019).

38. Stenum, T. S. et al. RNA interactome capture in Escherichia coli globally identifies RNA-binding proteins. Nucleic Acids Res. gkad216 (2023) doi:10.1093/nar/gkad216.

39. Szklarczyk, D. et al. The STRING database in 2023: protein–protein association networks and functional enrichment analyses for any sequenced genome of interest. Nucleic Acids Res. 51, D638–D646 (2023).

40. Kantrowitz, E. R. Allostery and cooperativity in *Escherichia coli* aspartate transcarbamoylase. Arch. Biochem. Biophys. 519, 81–90 (2012).

41. Sillitoe, I. et al. CATH: increased structural coverage of functional space. Nucleic Acids Res. 49, D266–D273 (2021).

42. Gutowski, S. J. & Rosenberg, H. Succinate uptake and related proton movements in Escherichia coli K12. Biochem. J. 152, 647–654 (1975).

43. Hart, M. L. et al. Succinate dehydrogenase loss suppresses pyrimidine biosynthesis via succinate-mediated inhibition of aspartate transcarbamylase. 2025.02.18.638948 Preprint at 10.1101/2025.02.18.638948 (2025).

44. Joyce, M. A. et al. Probing the Nucleotide-Binding Site of Escherichia coli Succinyl-CoA Synthetase. Biochemistry 38, 7273–7283 (1999).

45. Cockrell, G. M. et al. New Paradigm for Allosteric Regulation of Escherichia coli Aspartate Transcarbamoylase. Biochemistry 52, 8036–8047 (2013).

46. Huang, J. & Lipscomb, W. N. Products in the T-State of Aspartate Transcarbamylase: Crystal Structure of the Phosphate and N-Carbamyl-l-aspartate Ligated Enzyme,. Biochemistry 43, 6422–6426 (2004).

47. Kantrowitz, E. R. & Lipscomb, W. N. *Escherichia coli* aspartate transcarbamoylase: the molecular basis for a concerted allosteric transition. Trends Biochem. Sci. 15, 53–59 (1990).

48. Malinovska, L. et al. Proteome-wide structural changes measured with limited proteolysis-mass spectrometry: an advanced protocol for high-throughput applications. Nat. Protoc. 18, 659–682 (2023).

49. Lei, Z., Wang, N., Tan, H., Zheng, J. & Jia, Z. Conformational Plasticity of the Active Site Entrance in E. coli Aspartate Transcarbamoylase and Its Implication in Feedback Regulation. Int. J. Mol. Sci. 21, 320 (2020).

50. Welp, L. M. et al. Chemical crosslinking extends and complements UV crosslinking in analysis of RNA/DNA nucleic acid–protein interaction sites by mass spectrometry. Nucleic Acids Res. 53, gkaf727 (2025).

51. Begue, S. C., Leonardi, E. & Tosatto, S. C. E. Decoding protein structures with residue interaction networks. Trends Biochem. Sci. 50, 1072–1085 (2025).

52. Joyce, A. R. et al. Experimental and Computational Assessment of Conditionally Essential Genes in Escherichia coli. J. Bacteriol. 188, 8259–8271 (2006).

53. Burns, B. P. et al. The *Helicobacter pylori pyrB* Gene Encoding Aspartate Carbamoyltransferase Is Essential for Bacterial Survival. Arch. Biochem. Biophys. 380, 78–84 (2000).

54. Yang, H.-J., Bogomolnaya, L., McClelland, M. & Andrews-Polymenis, H. De novo pyrimidine synthesis is necessary for intestinal colonization of Salmonella Typhimurium in chicks. PLOS ONE 12, e0183751 (2017).

55. van Kempen, M. et al. Fast and accurate protein structure search with Foldseek. Nat. Biotechnol. 42, 243–246 (2024).

56. Liao, J.-Y. et al. RBPWorld for exploring functions and disease associations of RNA-binding proteins across species. Nucleic Acids Res. 53, D220–D232 (2025).

57. Leone, S. et al. HSP70 binds to specific non-coding RNA and regulates human RNA polymerase III. Mol. Cell 84, 687–701.e7 (2024).

58. Screen, M. et al. RNA helicase EIF4A1-mediated translation is essential for the GC response. Life Sci. Alliance 7, (2024).

59. Sfakianos, A. P. et al. Inhibiting translation elongation by reducing eIF5A activity induces feedback inhibition of initiation, limiting tumour cell proliferation. Nat. Commun. 16, 11486 (2025).

60. Kang, Y.-N., Tran, A., White, R. H. & Ealick, S. E. A Novel Function for the N-Terminal Nucleophile Hydrolase Fold Demonstrated by the Structure of an Archaeal Inosine Monophosphate Cyclohydrolase,. Biochemistry 46, 5050–5062 (2007).

61. De Donatis, D. et al. Duloxetine plasma level and antidepressant response. Prog. Neuropsychopharmacol. Biol. Psychiatry 92, 127–132 (2019).

62. Toda, T. et al. Protective Effects of Duloxetine against Cerebral Ischemia-Reperfusion Injury via Transient Receptor Potential Melastatin 2 Inhibition. J. Pharmacol. Exp. Ther. 368, 246–254 (2019).

63. Wang, M. et al. Duloxetine alleviates oxaliplatin-induced peripheral neuropathy by regulating p53-mediated apoptosis. NeuroReport 33, 437 (2022).

64. Hole, K. et al. Effect of CYP2D6 genotype on duloxetine serum concentration. Basic Clin. Pharmacol. Toxicol. 134, 186–192 (2024).

65. Zimova, L., Ptakova, A., Mitro, M., Krusek, J. & Vlachova, V. Activity dependent inhibition of TRPC1/4/5 channels by duloxetine involves voltage sensor-like domain. Biomed. Pharmacother. 152, 113262 (2022).

66. Urdaneta, E. C. et al. Purification of cross-linked RNA-protein complexes by phenol-toluol extraction. Nat. Commun. 10, 990 (2019).

67. Krause, K. L., Volz, K. W. & Lipscomb, W. N. Structure at 2.9-A resolution of aspartate carbamoyltransferase complexed with the bisubstrate analogue N-(phosphonacetyl)-L-aspartate. Proc. Natl. Acad. Sci. 82, 1643–1647 (1985).

68. Silver, R. S., Daigneault, J. P., Teague, P. D. & Kantrowitz, E. R. Analysis of two purified mutants of *Escherichia coli* aspartate transcarbamylase with single amino acid substitutions. J. Mol. Biol. 168, 729–745 (1983).

69. Horos, R. et al. The Small Non-coding Vault RNA1-1 Acts as a Riboregulator of Autophagy. Cell 176, 1054–1067.e12 (2019).

70. Miyakoshi, M., Matera, G., Maki, K., Sone, Y. & Vogel, J. Functional expansion of a TCA cycle operon mRNA by a 3′ end-derived small RNA. Nucleic Acids Res. 47, 2075–2088 (2019).

71. Castello, A. et al. Comprehensive Identification of RNA-Binding Domains in Human Cells. Mol. Cell 63, 696–710 (2016).

72. Hanukoglu, I. Proteopedia: Rossmann fold: A beta-alpha-beta fold at dinucleotide binding sites. Biochem. Mol. Biol. Educ. 43, 206–209 (2015).

73. Laurino, P. et al. An Ancient Fingerprint Indicates the Common Ancestry of Rossmann-Fold Enzymes Utilizing Different Ribose-Based Cofactors. PLOS Biol. 14, e1002396 (2016).

74. Rukavishnikov, G. et al. Antimicrobial activity of antidepressants on normal gut microbiota: Results of the in vitro study. Front. Behav. Neurosci. 17, (2023).

75. Li, Y. et al. Microbial metabolites in the gut-brain axis: their impact on depression pathophysiology and treatment. Neuroscience 593, 1–7 (2026).

76. Ohara, T. E. & Hsiao, E. Y. Microbiota–neuroepithelial signalling across the gut– brain axis. Nat. Rev. Microbiol. 23, 371–384 (2025).

77. Ritz, N. L., et al. Social anxiety disorder-associated gut microbiota increases social fear. Proc. Natl. Acad. Sci. 121, e2308706120 (2024).

78. Valles-Colomer, M. et al. The neuroactive potential of the human gut microbiota in quality of life and depression. Nat. Microbiol. 4, 623–632 (2019).

79. Wu, P. Inhibition of RNA-binding proteins with small molecules. Nat. Rev. Chem. 4, 441–458 (2020).

80. Jiang, X. et al. In-Depth Metaproteomics Analysis of Oral Microbiome for Lung Cancer. Research 2022, (2022).

81. Zhu, Y. et al. DEqMS: A Method for Accurate Variance Estimation in Differential Protein Expression Analysis *. Mol. Cell. Proteomics 19, 1047–1057 (2020).

82. Terpe, K. Overview of bacterial expression systems for heterologous protein production: from molecular and biochemical fundamentals to commercial systems. Appl. Microbiol. Biotechnol. 72, 211–222 (2006).

83. Pastra-Landis, S. C., Foote, J. & Kantrowitz, E. R. An improved colorimetric assay for aspartate and ornithine transcarbamylases. Anal. Biochem. 118, 358–363 (1981).

84. Lee, F. C. Y. et al. An improved iCLIP protocol. 2021.08.27.457890 Preprint at 10.1101/2021.08.27.457890 (2021).

85. Busch, A., Brüggemann, M., Ebersberger, S. & Zarnack, K. iCLIP data analysis: A complete pipeline from sequencing reads to RBP binding sites. Methods 178, 49– 62 (2020).

86. Robinson, J. T. et al. Integrative genomics viewer. Nat. Biotechnol. 29, 24–26 (2011).

87. Kuret, K., Amalietti, A. G., Jones, D. M., Capitanchik, C. & Ule, J. Positional motif analysis reveals the extent of specificity of protein-RNA interactions observed by CLIP. Genome Biol. 23, 191 (2022).

88. Bailey, T. L., Johnson, J., Grant, C. E. & Noble, W. S. The MEME Suite. Nucleic Acids Res. 43, W39–W49 (2015).

89. Lorenz, R. et al. ViennaRNA Package 2.0. Algorithms Mol. Biol. 6, 26 (2011).

90. Berman, H. M. et al. The Protein Data Bank. Nucleic Acids Res. 28, 235–242 (2000).

91. Jumper, J. et al. Highly accurate protein structure prediction with AlphaFold. Nature 596, 583–589 (2021).

92. Varadi, M. et al. AlphaFold Protein Structure Database in 2024: providing structure coverage for over 214 million protein sequences. Nucleic Acids Res. 52, D368– D375 (2024).

93. Meng, E. C. et al. UCSF ChimeraX: Tools for structure building and analysis. Protein Sci. 32, e4792 (2023).

94. Forli, S. et al. Computational protein–ligand docking and virtual drug screening with the AutoDock suite. Nat. Protoc. 11, 905–919 (2016).

95. Baek, M. et al. Accurate prediction of protein–nucleic acid complexes using RoseTTAFoldNA. Nat. Methods 21, 117–121 (2024).

96. Sarzynska, J., Popenda, M., Antczak, M. & Szachniuk, M. RNA tertiary structure prediction using RNAComposer in CASP15. Proteins Struct. Funct. Bioinforma. 91, 1790–1799 (2023).

97. Moafinejad, S. N. et al. SimRNAweb v2.0: a web server for RNA folding simulations and 3D structure modeling, with optional restraints and enhanced analysis of folding trajectories. Nucleic Acids Res. 52, W368–W373 (2024).

98. Giulini, M. et al. HADDOCK3: A Modular and Versatile Platform for Integrative Modeling of Biomolecular Complexes. J. Chem. Inf. Model. 65, 7315–7324 (2025).

99. Kagami, L., Wilter, A., Diaz, A. & Vranken, W. The ACPYPE web server for small-molecule MD topology generation. Bioinformatics 39, btad350 (2023).

100. Krissinel, E. & Henrick, K. Inference of Macromolecular Assemblies from Crystalline State. J. Mol. Biol. 372, 774–797 (2007).

101. Ono, K. et al. Cytoscape Web: bringing network biology to the browser. Nucleic Acids Res. 53, W203–W212 (2025).

102. O’Mahoney, C. et al. Microfluidic Diffusional Sizing (MDS) Measurements of Secretory Neutralizing Antibody Affinity Against SARS-CoV-2. Ann. Biomed. Eng. 52, 1653–1664 (2024).

103. Letunic, I. & Bork, P. Interactive Tree of Life (iTOL) v6: recent updates to the phylogenetic tree display and annotation tool. Nucleic Acids Res. 52, W78–W82 (2024).

