## Supporting figures for "Network-scale disruption of RNA-protein interactions by a bioaccumulating small-molecule drug"

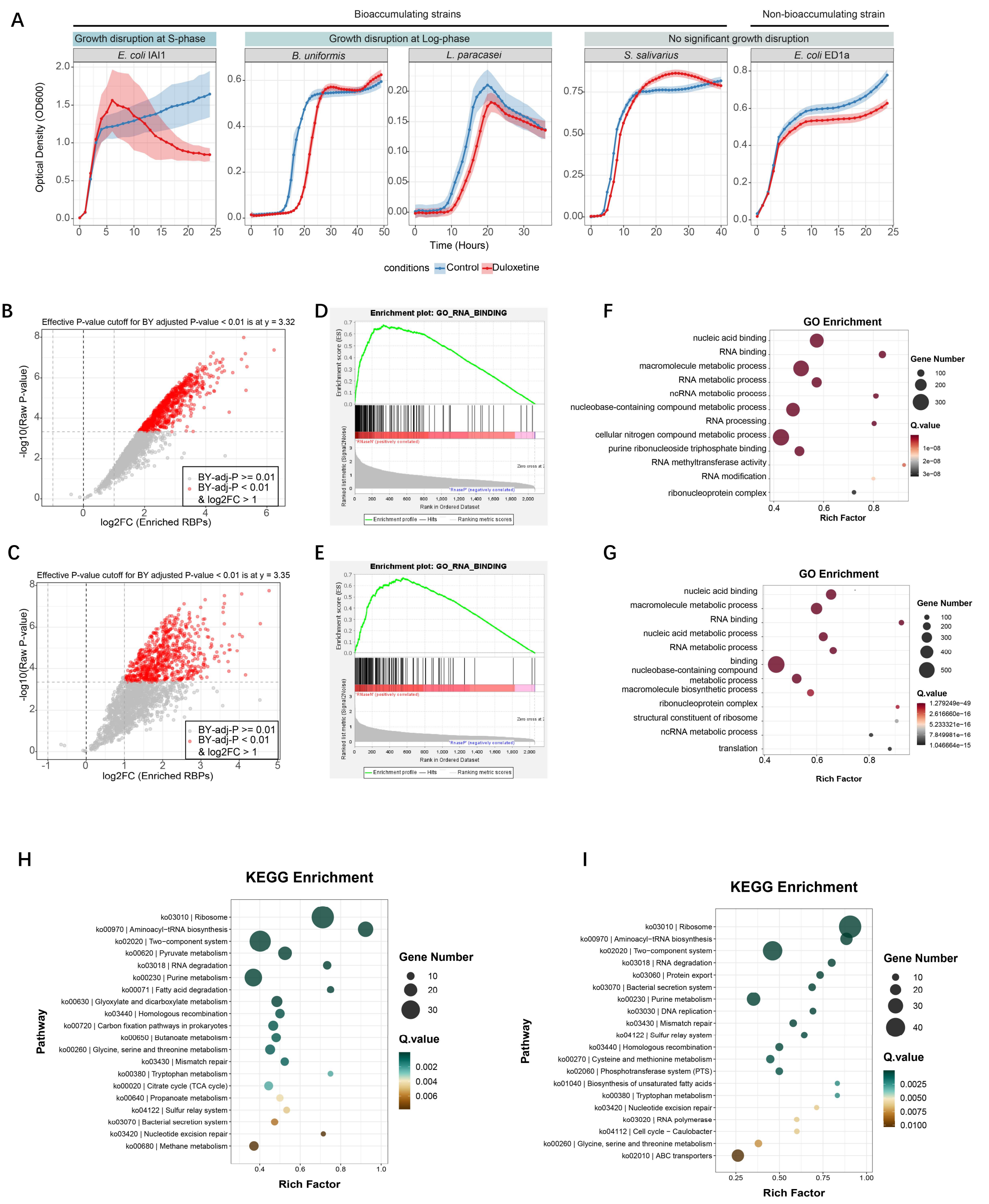


**Figure S1 Functional enrichment analysis of OOPS-enriched RBP candidates.**

**A,** Growth curve of four duloxetine-bioaccumulated strains and one non-bioaccumulated strain with/without duloxetine.

**B, C,** Volcano plot showing RNA-binding protein enrichment from *E. coli* IAI1 and ED1a, respectively.

**D, E,** Gene set enrichment analysis (GESA) demonstrating significant enrichment of the RNA binding (GO:0003723) protein set at the leading edge of the ranked RBP datasets (Normalized Enrichment Score = 2.79 for IAI1, and 2.76 for ED1a, FDR q-val < 0.001).

**F, G,** Gene Ontology enrichment analysis of the identified RBP candidates in IAI1 and ED1a showing overrepresentation of RNA-related functions and metabolic processes.

**H, I,** KEGG pathway enrichment of RBP candidates identified in *E. coli* IAI1 and ED1a.

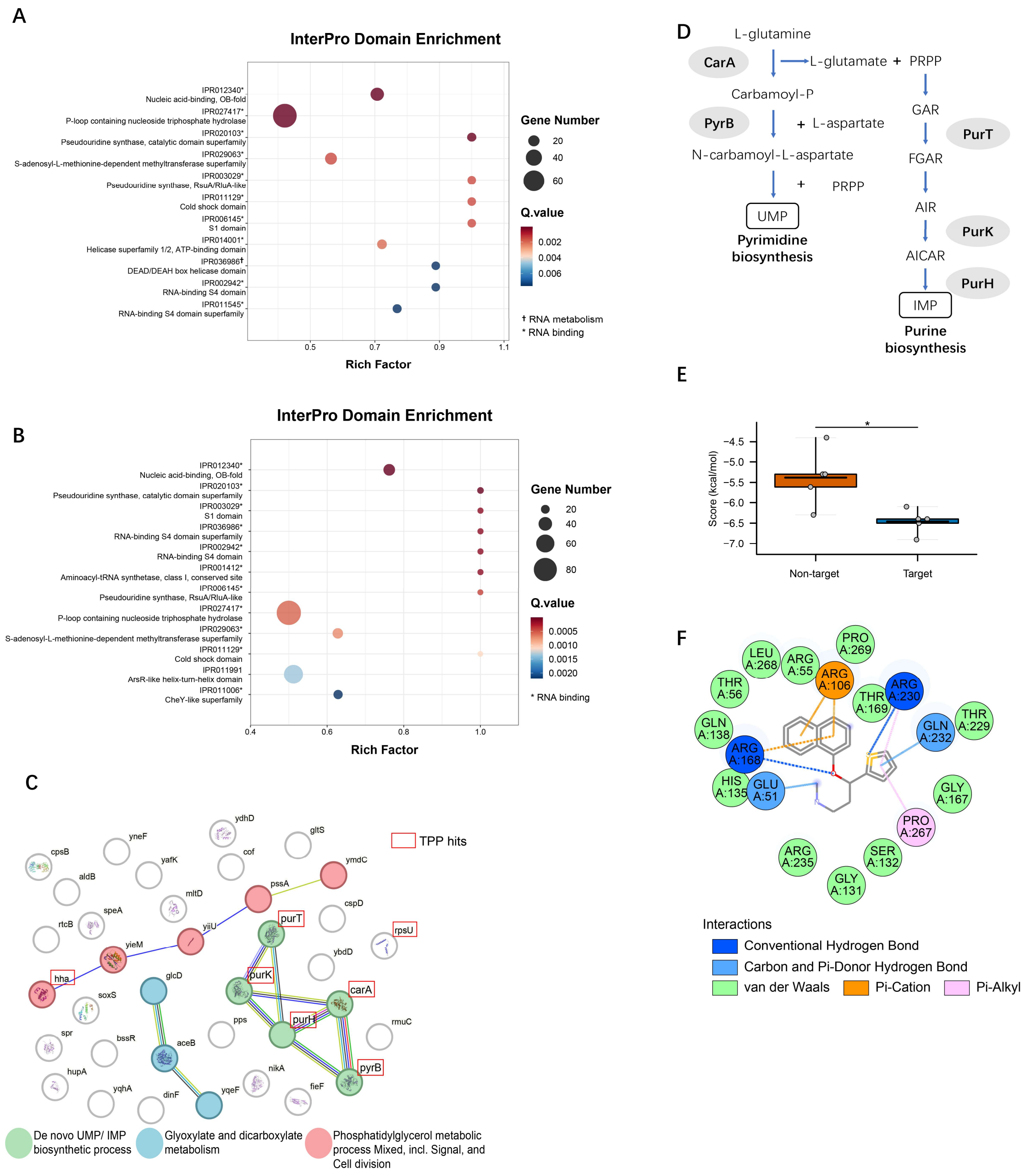


**Figure S2 Functional and structural characterization of duloxetine-disrupted RNA-binding proteins.**

**A, B,** InterPro Doman enrichment analysis of OOPS-enriched RBP candidates identified in *E. coli* IAI1 and ED1a.

**C,** STRING network analysis showing three major networks among the RNA-disrupted proteins by duloxetine. Red squares indicates the proteins overlap with the previous thermal proteome profiling (TPP) result (Klünemann et al., 2021), representing duloxetine-interacting proteins.

**D,** Metabolic network of the *de novo* UMP/IMP biosynthetic process identified in the STRING analysis.

**E,** Molecular docking scores of duloxetine against five key targets within the *de novo*

UMP/IMP biosynthetic network (n = 5) versus non-target proteins (n = 5). Non-targets were randomly selected from the unresponsive OOPS-DIA background, defined as proteins showing no significant alteration in RNA-binding capacity upon drug treatment. Lower docking scores indicate higher predicted binding affinity. Statistical significance was determined using an independent two-sided t-test (*$p < 0.05$).

**F,** 2D ligand-receptor interaction plot of duloxetine bound to PyrB derived from molecular docking.

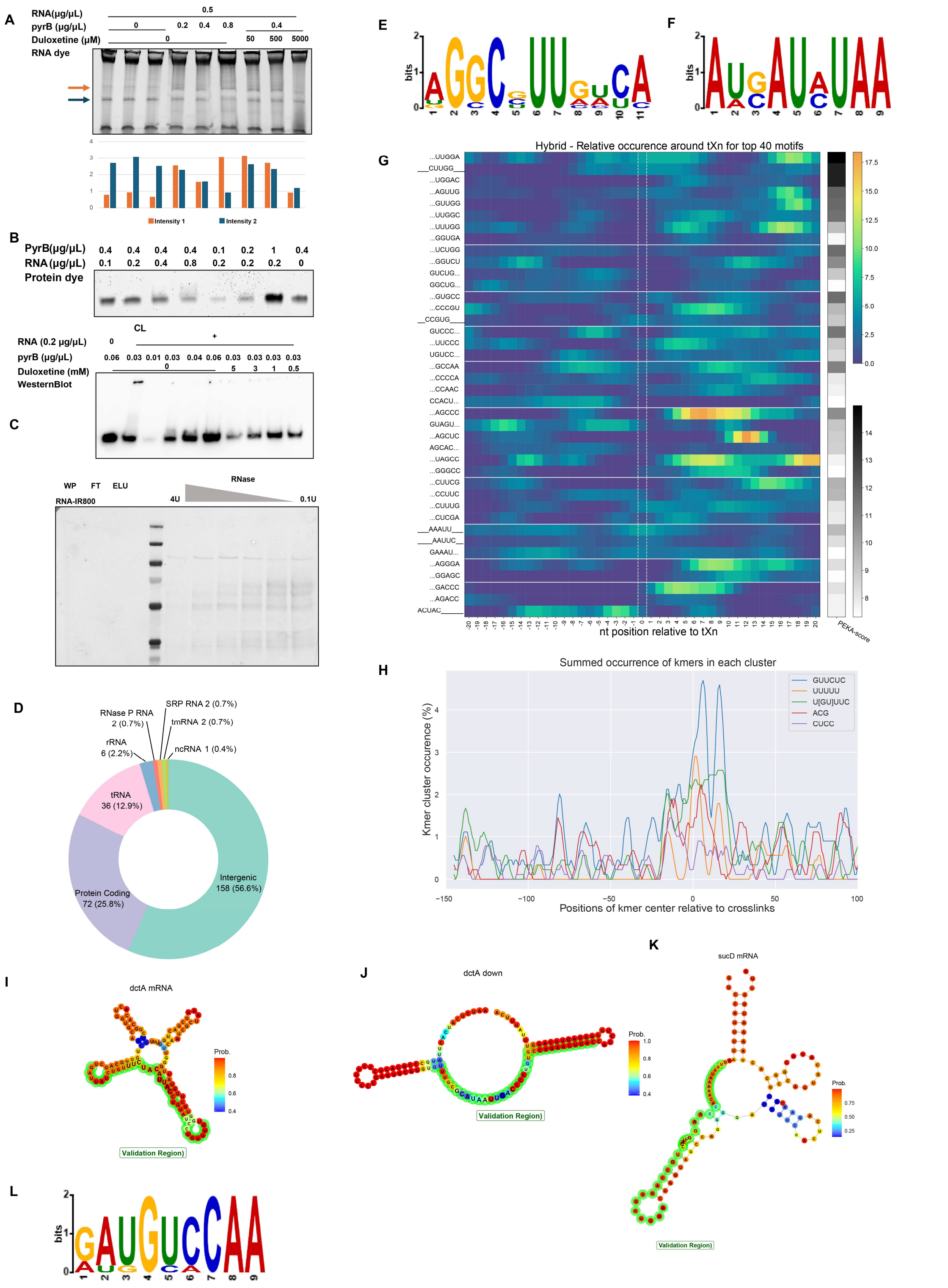

**Figure S3 PyrB recognizes different RNA motifs between intergenic and protein-coding regions.**

**A,** Top, electromobility shift assay (EMSA) of PyrB and the whole RNA extracted from the IAI1 strain. The gel was stained with SYBR Green to show the RNA pattern. The orange and dark blue arrows indicate the major RNA band shift. Bottom, the RNA band intensities extracted by imageJ. The colors correspond to the colors of arrows above.

**B,** Electrophoretic mobility shift assay (EMSA) of PyrB with total RNA extracted from the IAI1 strain. Both protein and RNA were used without labelling. Up plot, the gel was stained with SYPRO Ruby, showing a mobility shift corresponding to PyrB-RNA complex formation. The concentrations of RNA and protein are indicated in the figure. Bottom plot, Western blot analysis of the EMSA further confirms PyrB-RNA interactions. CL, UV cross-linked PyrB-RNA complex. Addition of RNA resulted in a smeared signal above the free protein band. Increasing concentrations of duloxetine led to PyrB precipitation and a reduced protein signal.

**C,** Visualization of biotin-labeled RNA on the nitrocellulose membrane. WP, whole protein without Ni-NTA magnetic bead enrichment. FT, flowthrough proteins after Ni-NTA magnetic bead enrichment. ELU, eluted protein after Ni-NTA magnetic bead enrichment. RNase gradients used in this experiment are listed on the plot.

**D,** PyrB interacting RNA classification.

**E,** RNA motif of the intergenic regions that PyrB interacts with, analysed by MEME (Bailey et al., 2015).

**F,** RNA motif of the protein-coding regions that PyrB interacts with.

**G,** Heatmap of relative occurrences (RtXn) and positionally enriched k-mer analysis (PEKA) scores for top 40 k-mers identified from PyrB-interacting intergenic RNA sequences. Three dots representing the maximal occurrence position is located further than 3 nt from the crosslink site.

**H,** PEKA analysis showing the frequency of PyrB-interacting protein-coding RNA motifs and their positions.

**I, J, K,** RNA structure of the PyrB targets *dctA* mRNA, *dctA* RNA 3'UTR, and *sucD* mRNA predicted by ViennaRNA. The highlighted regions (35 nt) were expanded from the 11 nt PyrB-interacting RNA regions. Enlarged nucleotide letters represent the 11 nt binding sites.

**L,** RNA motif from PyrB binding sequences.

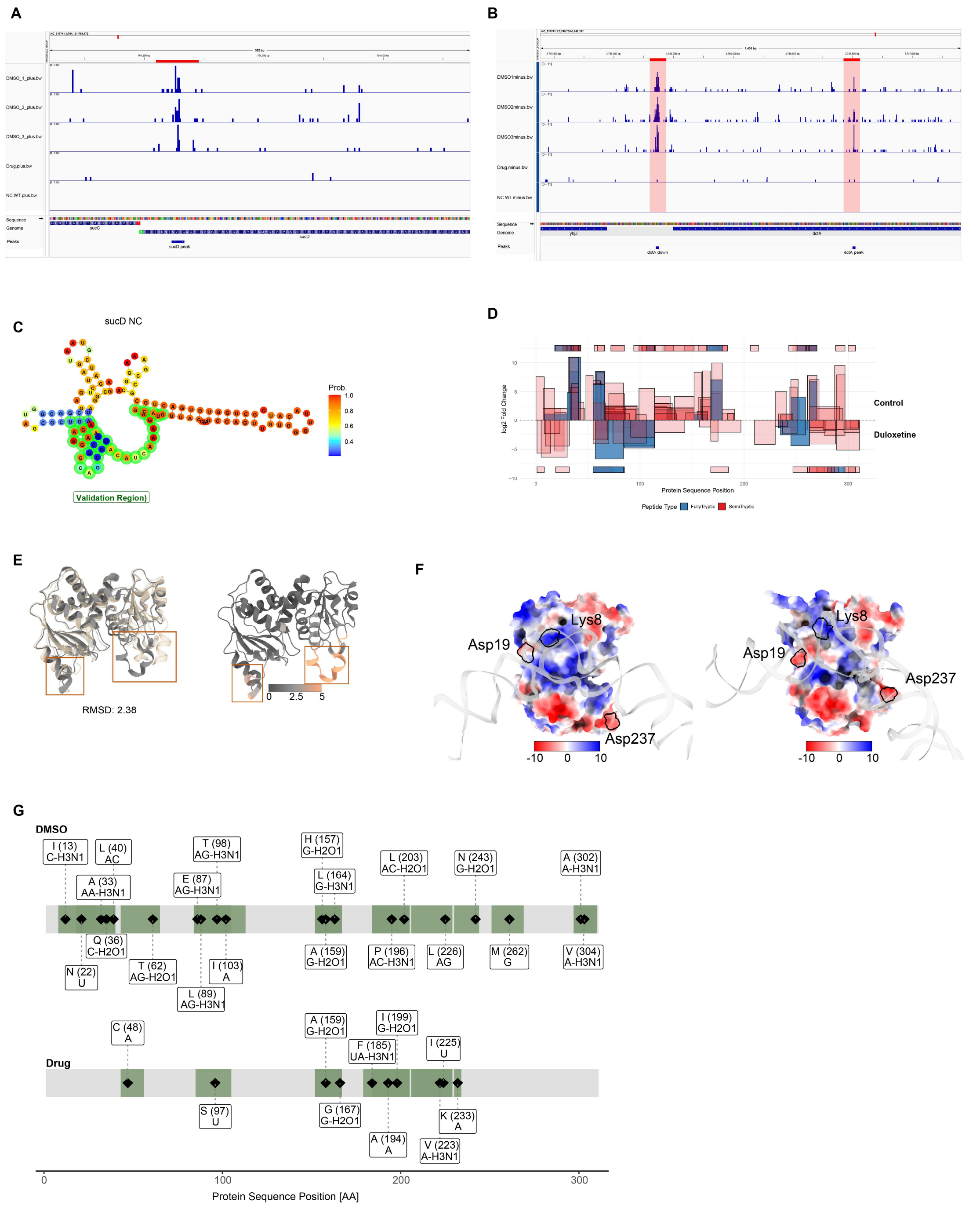


**Figure S4 Supplementary structural and biochemical characterization of PyrB-RNA interactions and duloxetine-induced conformational remodeling.**

**A, B,** iCLIP signal tracks of the 3 replicates DMSO control, the merged signal of 3 duloxetine replicates, and the wild type negative control on the *sucD* mRNA and the *dctA* RNA.

**C,** Secondary structure prediction of the *sucD* negative control (NC) RNA region generated by ViennaRNA. The highlighted 35-nucleotide validation region was selected for RNA synthesis.

**D,** Mapping of LiP-MS identified peptides and their corresponding log2 fold changes (duloxetine treatment vs. DMSO control) across the PyrB protein sequence. Fully tryptic peptides are colored in blue, and semi-tryptic peptides are in red. Infinite (INF) fold changes, representing peptides exclusively detected in one condition but completely absent in the other, are plotted at the maximum/minimum y-axis margins. All displayed peptides are filtered at $P < 0.01$.

**E,** Structural comparison between the active and inactive states of PyrB. **Left:** Structural overlay of the active (dark gray) and inactive (beige) PyrB conformations (overall RMSD: 2.38 Å). **Right:** The active state PyrB mapped with RMSD scores to highlight regions of high structural deviation relative to the inactive state.

**F,** Electrostatic surface potential of PyrB during *sucD* 3' UTR interaction (**left**) and following duloxetine binding (**right**). Positively and negatively charged regions are shown in blue and red, respectively (scale: -10 to +10). Black outlines highlight three representative regions exhibiting significant electrostatic redistribution upon drug binding. The light grey backbone represents the modeled RNA.

**G,** Identification of RNA-interacting peptides on PyrB using RNA-XL-MS. The sequence maps display crosslinked peptides for the DMSO control (**top**) and duloxetine-treated samples (**bottom**). Green shaded boxes denote identified RNA-interacting peptide regions. Black diamonds indicate specific crosslinked amino acid residues, annotated with their corresponding positional numbers and nucleotide adduct mass shifts.

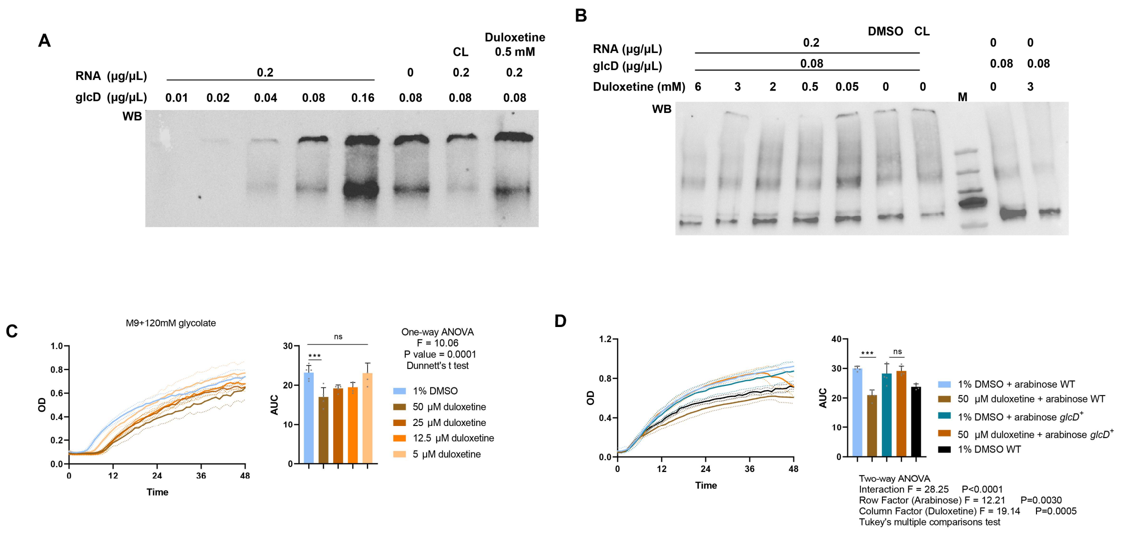


**Figure S5 Duloxetine can interact with another RBP, GlcD, disrupting its enzyme function.**

**A,** Western blot of the electromobility shift assay (EMSA) of different concentrations of GlcD and the whole RNA extracted from the IAI1 strain. CL, UV cross-linked PyrB-RNA complex.

**B,** Western blot of the electromobility shift assay (EMSA) of GlcD and the whole RNA extracted from the IAI1 strain with different concentrations of duloxetine. CL, UV cross-linked PyrB-RNA complex.

**C,** Growth curves and area under the curve values of *E. coli* IAI1 on M9+120 mM glycolate media with different duloxetine concentrations. The effect of growth inhibition was calculated using one-way ANOVA (F = 10.06, P = 0.0001, Dunnett's t test). WT, wild type.

**D,** Growth curves and area under the curve values of *E. coli* IAI1 wild type (WT) and GlcD overexpression IAI1 strain (*glcD*$^+$, 0.01% arabinose as inducer) on M9+120 mM glycolate media. The effect of growth inhibition was calculated using two-way ANOVA, with the interaction F = 28.25, P < 0.0001, row factor (arabinose) F = 12.21, P = 0.003, column factor (duloxetine) F = 19.14, P = 0.0005, Tukey's multiple comparisons test.

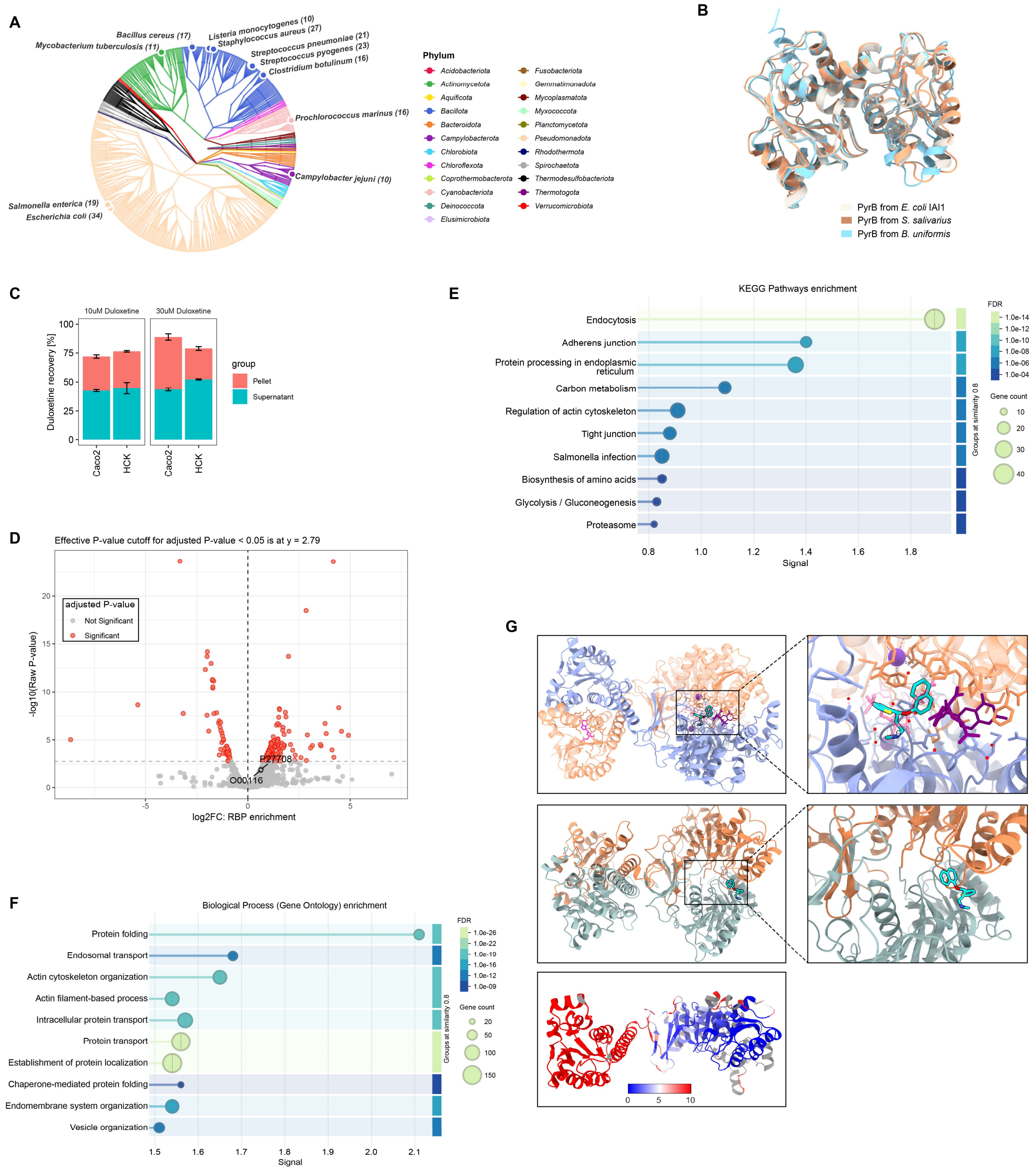

**Figure S6 Human cells bioaccumulate duloxetine, leading to disrupted RNA-protein interactions**

**A,** Taxonomic distribution of high-confidence structural homologs of *E. coli* PyrB. The phylogenetic tree illustrates the widespread conservation of this structural fold across diverse bacterial phyla.

**B,** Structural superimposition of AlphaFold-predicted PyrB orthologs from *B. uniformis* (blue) and *S. salivarius* (pink) against the *E. coli* IAI1 PyrB reference structure (light gray). Alignment of the core fold reveals highly conserved structural topologies across species. For *B. uniformis*, the pruned RMSD is 0.792 Å over 264 Cα atom pairs (global RMSD = 1.545 Å across 302 aligned pairs). For *S. salivarius*, the pruned RMSD is 0.942 Å over 217 core Cα atom pairs (global RMSD = 3.065 Å across 293 aligned pairs).

**C,** Quantification of duloxetine bioaccumulation in human intestinal epithelial (Caco-2) and renal (HCK) cell lines. The bar charts show the percentage of duloxetine recovered from the cellular pellet versus the supernatant following 10 μM and 30 μM treatments.

**D,** Volcano plot depicting the RNA-binding protein enrichment from human Caco-2 cells under duloxetine treatment. Enrichment is determined by comparing the RNase-negative experimental group against the RNase-positive control.

**E, F,** Functional characterization of duloxetine-disrupted RNA-binding proteins in Caco-2 cells. Bubble plots show the significant enrichment of (D) KEGG pathways and (E) Gene Ontology (GO) Biological Processes. Node size corresponds to the number of genes, and the color gradient represents the False Discovery Rate (FDR).

**G,** The structural similarity of human and *E. coli* AICAR transformylase domain. Top, the human ATIC protein structure (PDB 1P4R) and molecular docking result of duloxetine. Protein chains A and B are colored light purple and salmon, respectively. Endogenous ligands include 10-dideaza-folate (dark purple), AICAR (5-aminoimidazole-4-carboxamide ribonucleotide, pink), a potassium ion (purple sphere), and XMP (Xanthosine-5'-monophosphate, magenta). Duloxetine is highlighted in cyan with a black outline. Left plot, side view of the protein complex. The zoomed-in view (right) illustrates duloxetine competitively occupying the AICAR binding pocket. Middle, molecular docking of duloxetine into the *E. coli* PurH protein structure (PDB: 3ZZM). Protein chains A and B are colored light blue and salmon, respectively. The zoomed-in view (right) maps the duloxetine binding position within the catalytic cleft. Bottom, structural alignment between human ATIC (PDB 1P4R) and *E. coli* PurH (PDB 3ZZM). The color gradient from blue to red maps the Root-Mean-Square Deviation (RMSD) scores. For visual clarity, only a single chain of 1P4R is displayed in the alignment.

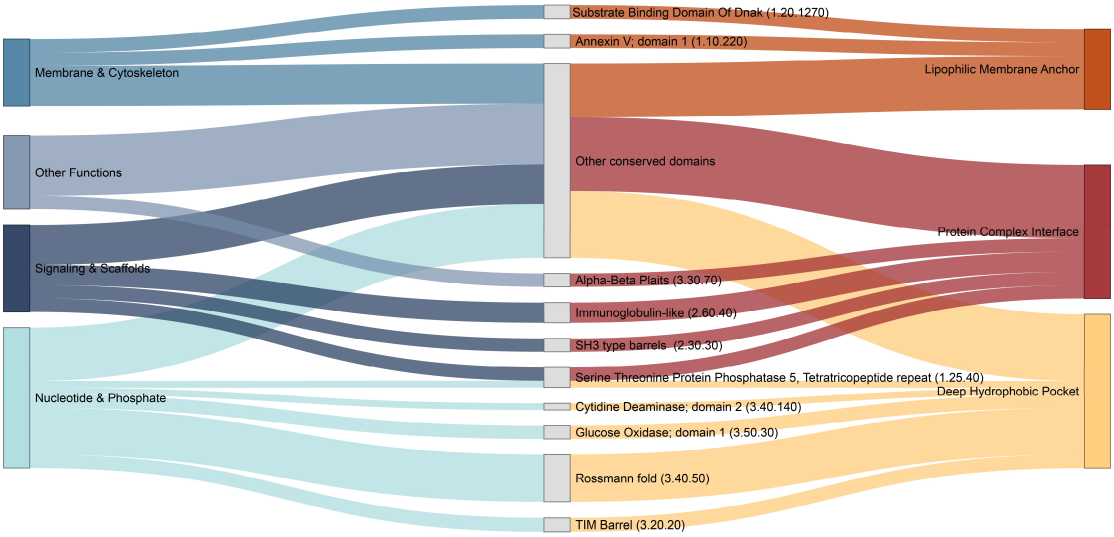


**Figure S7. Duloxetine targets conserved structural/functional pockets shared across bacterial and human proteomes.** Sankey diagram illustrating the mechanistic connections between primary biological functions (left column), enriched CATH structural domains (middle column), and predicted structural modes of action (right column) for duloxetine-disrupted RBPs in both *E. coli* IAI1 and human Caco-2 cells. The flow visualization demonstrates how functionally diverse proteins funnel into specific, shared physical vulnerabilities exploited by the drug. The structural modes of action on the right are categorized using combinations of four primary structural features: A, Hydrophobic Pocket; B, Protein Complex (interaction interfaces); C, Conformational Switch; and D, Membrane-Associated regions.
